# Activation of the bridge-like lipid transfer protein Atg2 by Atg1-mediated phosphorylation

**DOI:** 10.64898/2026.05.15.725374

**Authors:** Sonja Achleitner, Susanna Tulli, Ainara Claveras Cabezudo, Martina Schuschnig, Verena Baumann, Maximilian Schmid, Gerhard Hummer, Sascha Martens

## Abstract

Bridge-like lipid transfer proteins (BLTPs) have emerged as major contributors to lipid flux and thus cellular organization. BLTPs harbor a hydrophobic channel connecting two membranes to enable the bulk flow of lipids between them. The regulation of BLTPs is incompletely understood. Employing *in vitro* reconstitution, molecular dynamics simulations, and cell biology, we discovered that the BLTP Atg2, which mediates lipid transfer during autophagosome biogenesis, is activated by the Atg1 kinase. Atg1 phosphorylates two serines in the N-terminal region of Atg2. This triggers membrane binding and is predicted to open the channel for lipid transfer. The Atg1 kinase complex is localized to the endoplasmic reticulum in part through binding of its Atg13 subunit to the VAP protein Scs2. Phosphorylated Atg2 can subsequently establish contacts with Atg9 vesicles. Our study delineates a pathway for the Atg1 kinase mediated activation of the BLTP Atg2 to promote membrane contact site formation and lipid flux.

## Introduction

Membrane contact sites (MCSs) are regions of close contact between cellular membrane-bound compartments. MCSs have emerged as major contributors to cellular organization by mediating flow of substances and metabolites including lipids^1–3^. Recent estimates suggest that non-vesicular transport of lipids at MCSs accounts for a large fraction of lipid flux within cells (Iglesias-Artola *et al*, 2025). Lipid transfer at MCSs can be mediated by shuttle-like lipid transfer proteins, which move individual lipids between compartments (Reinisch *et al*, 2025; Prinz *et al*, 2020). Apart from these shuttles, bridge-like lipid transfer proteins (BLTPs) have arisen as highways for the bulk lipid transfer between separate membranes. The BLTPs span distances of about 10-30 nm and allow lipid flux through a hydrophobic channel or groove spanning the entire length of the molecule connecting the two membranes (Levine, 2022). Lipid transfer proteins at MCSs are frequently recruited by VAMP-associated proteins (VAPs) in the ER, which bind to FFAT (two phenylalanines in an acidic tract) motifs and which are often regulated by phosphorylation (Levine, 2025).

Among the most extensively studied BLTPs is Atg2, which localizes to the MCS between the endoplasmic reticulum (ER) and the phagophore during autophagosome biogenesis. Autophagosome biogenesis requires the mobilization of millions of lipids on the timescale of minutes, and Atg2 is a major contributor to this lipid flux from the ER into the expanding phagophore membrane (Kotani *et al*, 2018; Osawa *et al*, 2019; Maeda *et al*, 2019; Valverde *et al*, 2019).

The seed for the phagophore is provided, at least in part, by Golgi-derived vesicles containing the transmembrane protein Atg9 (Yamamoto *et al*, 2012; Sawa-Makarska *et al*, 2020; Olivas *et al*, 2023). These seeds are connected to the ER via Atg2 and convert into the growing phagophore as they receive lipids from the ER via Atg2 (Holzer *et al*, 2024). Atg9 directly interacts with the Atg2 C-terminus and its lipid scramblase activity distributes the incoming lipids to both leaflets of the phagophore allowing for membrane growth (Maeda *et al*, 2020; Matoba *et al*, 2020; Ghanbarpour *et al*, 2021; Gómez-Sánchez *et al*, 2018; Van Vliet *et al*, 2022). The driving force for this net transfer towards the growing phagophore may arise from newly synthesized lipids in the ER close to the site of phagophore growth (Nishimura *et al*, 2017; Andrejeva *et al*, 2020; Schütter *et al*, 2020; Kannan *et al*, 2017; Nebauer *et al*, 2007; Rockenfeller *et al*, 2015; Baumann *et al*, 2024).

Autophagy is initiated by the assembly of the Atg1 kinase complex consisting of the conserved serine/threonine kinase Atg1 and Atg13 (Papinski & Kraft, 2016). In yeast, the Atg1 kinase complex also contains the Atg17-Atg31-Atg29 subcomplex (Stjepanovic *et al*, 2014; Suzuki *et al*, 2001). Following Atg1 recruitment and activation, components of the autophagy machinery assemble at the phagophore assembly site. Among the known targets of Atg1 kinase activity are Atg2, Atg4, and Atg9 (Fujioka *et al*, 2014; Yamamoto *et al*, 2016; Matsuura *et al*, 1997; Papinski *et al*, 2014). While the phosphorylation of Atg9 and Atg4 are important for the recruitment of downstream factors and inhibition of premature Atg8 deconjugation, respectively, the function of phosphorylation of the BLTP Atg2 is still not resolved (Papinski *et al*, 2014; Sánchez-Wandelmer *et al*, 2017; Hu *et al*, 2019).

We set out to shed light on the spatiotemporal regulation of Atg2 at the ER-phagophore MCS. Employing *in vitro* reconstitution, molecular dynamics (MD) simulations as well as cell biology we found that Atg2 is phosphorylated by Atg1 at its N-terminus. This phosphorylation triggers a conformational change, resulting in the displacement of helix 1 (H1) and helix 2 (H2), enhancing membrane binding, and opening the groove for lipid transfer. Further, we found that the ER-phagophore MCS is coordinated by the ER-residing VAP family protein Scs2 by interacting with the Atg1 complex subunits Atg13 and Atg1 as well as Atg2 in a phospho-regulated manner. Our study reveals a phosphorylation mediated activation of Atg2 with implications for BLTP activation in general.

## Results

### Atg1 stimulates Atg2-mediated lipid transfer by phosphorylation

The Atg1 kinase complex is a master regulator of autophagy initiation and progression. We therefore asked if it has a direct effect on the lipid transfer activity of Atg2. To this end, we employed an *in vitro* Förster resonance energy transfer (FRET)-based lipid transfer assay, which is based on the de-quenching of the NBD fluorophore (Osawa *et al*, 2019) (Fig. 1A). Atg2 was purified in complex with Atg18 because co-purification consistently showed increased Atg2 stability compared to Atg2 purification alone. In addition, we purified the Atg1 kinase and Atg13, which represent the evolutionary conserved subunits of the Atg1/ULK1 complex (Papinski & Kraft, 2016). Interestingly, we observed a robust stimulation of lipid transfer by Atg2 in presence of Atg1-Atg13 and ATP but not the non-hydrolysable analogue AMP-PNP (Fig. 1B). This suggested that the Atg1 kinase activity is required for the increase in Atg2 activity. Consistently, the Atg1 kinase alone was sufficient to mediate the ATP-dependent stimulation of lipid transfer (Fig. 1C). Next, we asked if the lipid transfer enhancing effect was exclusively mediated by phosphorylation of the Atg2-Atg18 complex or influenced by the presence of Atg1 in a kinase-independent manner as well. To distinguish between these possibilities, we preincubated the Atg2-Atg18 complex with Atg1 in the presence and absence of ATP, followed by purification of Atg2-Atg18 by size exclusion chromatography to remove Atg1. Although pre-phosphorylated Atg2-Atg18 showed a lower lipid transfer maximum than Atg2-Atg18 in the presence of Atg1, presumably due to some loss of activity during the lengthy procedure, the lipid transfer rate, estimated based on the time at which the half maximum was reached, was comparable (Fig. 1D). This indicated that Atg1 presence *per se* does not influence transfer of Atg2-Atg18. Additionally, we performed the FRET-based lipid transfer assay with a variant of Atg1 that only contained the kinase domain and excluded its membrane binding EAT domain (Atg1^1–454^) (Ragusa *et al*, 2012). This variant still stimulated the transfer activity of Atg2-Atg18 (Fig. 1E). Atg1 also stimulated the lipid transfer in the absence of Atg18 showing that phosphorylation of Atg2 is sufficient for this effect (Fig. EV1A).

**Figure 1.**
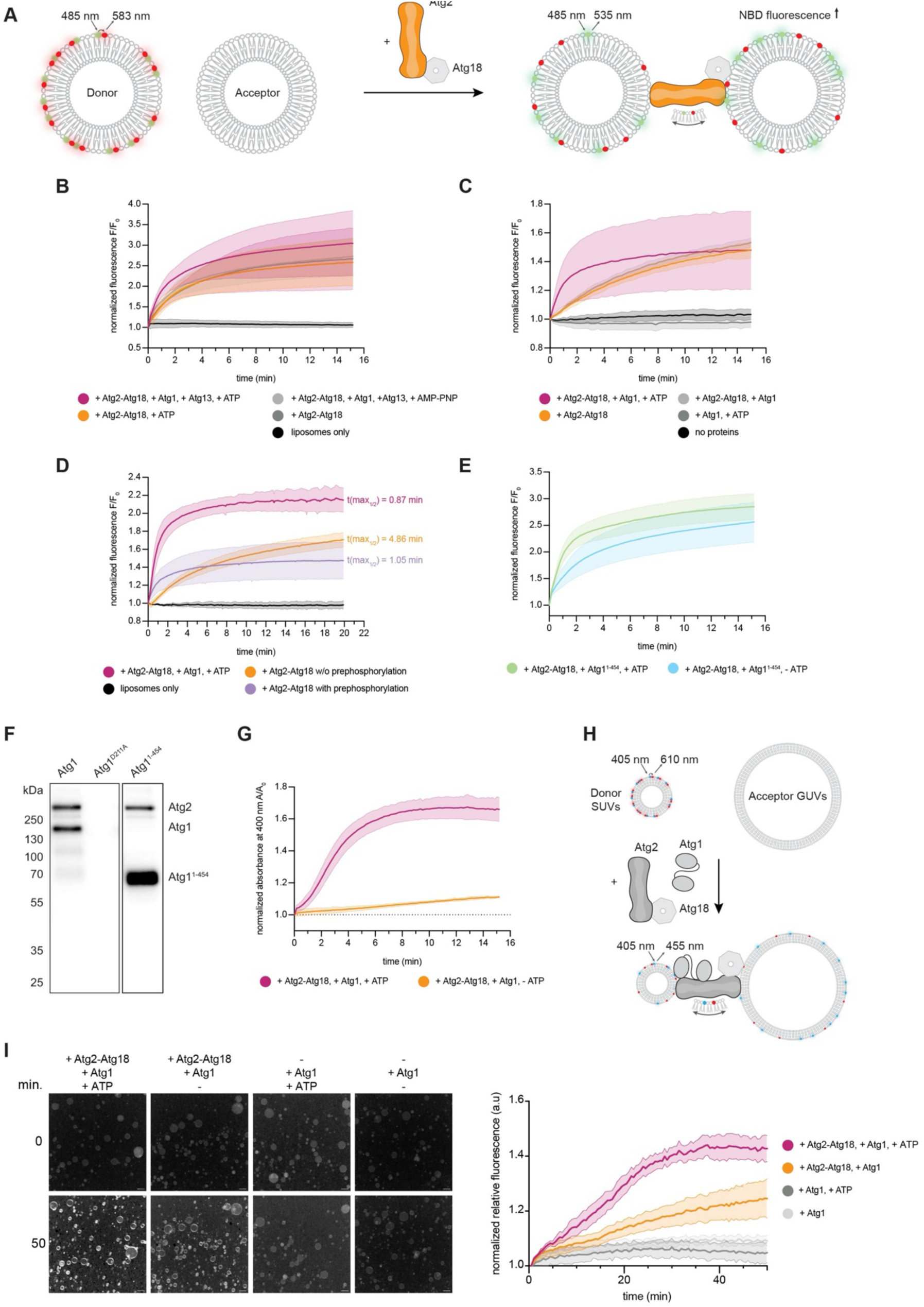
Atg1-dependent phosphorylation of Atg2 promotes lipid transfer. (**A**) Scheme of the FRET-based lipid transfer assay. Fluorescence of NBD-PE increases upon dilution into the acceptor membranes indicating transfer via Atg2. (**B**) FRET-based lipid transfer assay with ER-like and Atg9-vesicle-like liposomes showing that Atg1-Atg13 boosts Atg2-Atg18 lipid transfer in a kinase-dependent manner. (**C**) FRET-based lipid transfer assay showing Atg1 is sufficient to boost Atg2-Atg18 mediated lipid transfer in a kinase activity dependent manner. (**D**) FRET-based lipid transfer assay comparing pre-phosphorylated, non-pre-phosphorylated and Atg2-Atg18 including active Atg1. Pre-phosphorylated Atg2-Atg18 shows a similar time of half maximum fluorescence as Atg2-Atg18 in the presence of Atg1. (**E**) Atg1^1-454^ can still boost the lipid transfer by Atg2 as tested with the FRET-based transfer assay. (**F**) Radioactive kinase assay showing that Atg1 as well as Atg1^1-454^ can autophosphorylate and phosphorylate Atg2. (**G**) Absorbance-based turbidity assay to test Atg2 membrane tethering activity. Donor liposomes for b-g contained 66% DOPC, 10% DOPS, 20% DOPE, 2% lissamine rhodamine-DHPE, and 2% NBD-DPPE and the acceptor membranes contained 44% POPC, 6% POPS, 6% POPE, 41.5% liver PI, and 2.5% PI3P. The data are means ± SD (n = 3). (**H**) Scheme of the microscopy-based lipid transfer assay with GUVs and liposomes. Fluorescence of ATTO 425 increases upon dilution after Atg2-mediated lipid transfer. Atg2 tethered donor liposomes (N-terminus) and PI3P-containing GUVs (C-terminus). The ratio between the fluorescence of the quenched fluorophore (ATTO 425) and the quencher (ATTO 540Ǫ) accounted for lipid transfer. (**I**) Microscopy-based lipid transfer assay with GUVs and liposomes showing that Atg1 boosts Atg2-Atg18 mediated lipid transfer in a kinase activity dependent manner. Left, representative images showing ATTO 425/ATTO 540Ǫ ratio signal after 0 and 50 minutes of acquisition in described experimental conditions. Right, quantification of ATTO 425/ATTO 540Ǫ ratio signal over time. Multiple (5<x<30) ROIs per condition. Data are means ± SEM (n≥ 3), scale bars, 100 µm. Donor liposomes contained 66% DOPC, 10% DOPS, 20% DOPE, 2% ATTO 425 DOPE, and 2% ATTO 540Ǫ DOPE and the acceptor GUVs contained 75% DOPC, 10% DOPS, 10% DOPE, and 5% PI3P.

Collectively, these data strongly suggest that Atg1 phosphorylates Atg2, causing an increase in lipid transfer. This was confirmed in an *in vitro* phosphorylation assay, in which incubation of Atg2 with full length Atg1 or the kinase domain of Atg1 (1-454), but not with a kinase-deficient (D211A) version of Atg1, resulted in Atg2 phosphorylation (Fig. 1F). Next, we used a turbidity-based assay to test Atg2 membrane tethering. The phosphorylation-dependent boost of lipid transfer by the Atg2-Atg18 complex correlated with an increase in membrane tethering (Fig. 1G). We conclude that Atg1 enhances lipid transfer by Atg2 in a phosphorylation-dependent manner, likely in part by increasing membrane binding and thus tethering by Atg2.

We observed that NBD-fluorescence slightly increases in the FRET-based transfer assay also in the absence of acceptor liposomes indicating that dequenching already occurs upon the entry of lipids into the Atg2 cavity. However, the increase in fluorescence is much stronger in the presence of acceptor membranes suggesting that dequenching observed is mainly the result of lipid transfer (Fig. EV1B).

To corroborate the Atg1-mediated boost of the Atg2 lipid transfer activity, we established a microscopy-based FRET-assay with giant unilamellar vesicles (GUVs) and liposomes. Here, Atg2 was tethered to PI3P-containing GUVs via its C-terminus through the interaction between Atg18 and PI3P. Liposomes containing a FRET-pair (ATTO 425 and ATTO 540Ǫ) serve as the donor membranes (Holzer *et al*, 2026) (Fig. 1H). The ratio between the fluorescence of the quenched fluorophore (ATTO 425) and the quencher (ATTO 540Ǫ) accounted for lipid transfer. In the presence of Atg2-Atg18 and Atg1 ATTO 425 fluorescence increased over time indicating transfer of lipids. In line with the FRET-based assay with only liposomes, the addition of ATP also boosted lipid transfer between liposomes and GUVs (Fig. 1I).

### Phosphorylation of the Atg2 N-terminus enhances lipid transfer

To identify the Atg1 phosphorylation sites in Atg2 required for lipid transfer stimulation, we subjected the *in vitro* phosphorylated Atg2-Atg18 complex to mass spectrometry. Based on this analysis we identified 43 phospho-sites in Atg2, distributed throughout the entire protein (labelled red in Fig. 2A, Table 1, Dataset EV1). Among those sites were serines 98, 249, 1086 and 1113, which were previously reported (Papinski *et al*, 2014; Hu *et al*, 2019). To test if any of the 43 sites are required for the Atg1-dependent lipid transfer boost, we purified a Atg2-Atg18 version where the identified sites were mutated to alanine (Atg2^43xA^-Atg18). The basal lipid transfer activity of the Atg2^43xA^-Atg18 was low, suggesting partial destabilization of the mutant Atg2. No increase in lipid transfer by Atg1 was detectable (Fig. EV1C). To prioritize the phosphorylation sites to be further tested, we reasoned that the relevant phospho-sites are likely located in the N- and/or C-terminal regions of Atg2, because these regions mediate Atg2 membrane binding. We therefore generated two Atg2 mutants: one with ten phospho-sites in the N-terminal region mutated to alanines (Atg2^N-10xA^-Atg18; Table 1) and one with twelve phospho-sites in the C-terminal region replaced with alanines (Atg2^C-12xA^-Atg18; Table 1). In the *in vitro* lipid transfer assay, the Atg2^N-10xA^-Atg18 mutant showed reduced transfer stimulation by Atg1, while the Atg2^C-12xA^-Atg18 behaved similar to the wild type protein (Fig. EV1D,E). We concluded that the relevant Atg1 target sites reside in the N-terminal region of Atg2. AlphaFold3 predictions indicated that the mutated sites in the Atg2^N-10xA^-Atg18 construct lie within helix H2 (residues 104-118, orange in Fig. 2A) and adjacent regions. Interestingly, one of these residues, serine at position 116 (red in zoom in Fig. 2A), corresponds to the Atg1 phosphorylation consensus sequence (Papinski *et al*, 2014; Hu *et al*, 2019). Additionally, we found that serine 109 (blue in zoom in Fig. 2A), which was not included in the Atg2^43xA^ mutant, also conforms to the Atg1 consensus sequence. While this site was not identified in the initial mass spectrometry analysis, digestion with chymotrypsin additionally to trypsin, led to the detection of peptides containing S109ph (Dataset EV1). Mass spectrometry analysis of Atg2-3xGFP isolated from yeast cells revealed that these two sites are also present in vivo upon induction of autophagy by nitrogen starvation and only in cells expressing Atg1 (Fig. 2B, Appendix Fig. S1). Mutating either of these two sites mimicked the Atg2^N-10xA^ mutant in the *in vitro* transfer assay. The transfer was reduced in both mutants and stimulation by Atg1 was impaired (Fig. EV1F,G). Interestingly, transfer by the double-mutant Atg2^S109A,S116A^ was almost abolished and could not be enhanced by Atg1 kinase activity (Fig. 2C).

**Figure 2.**
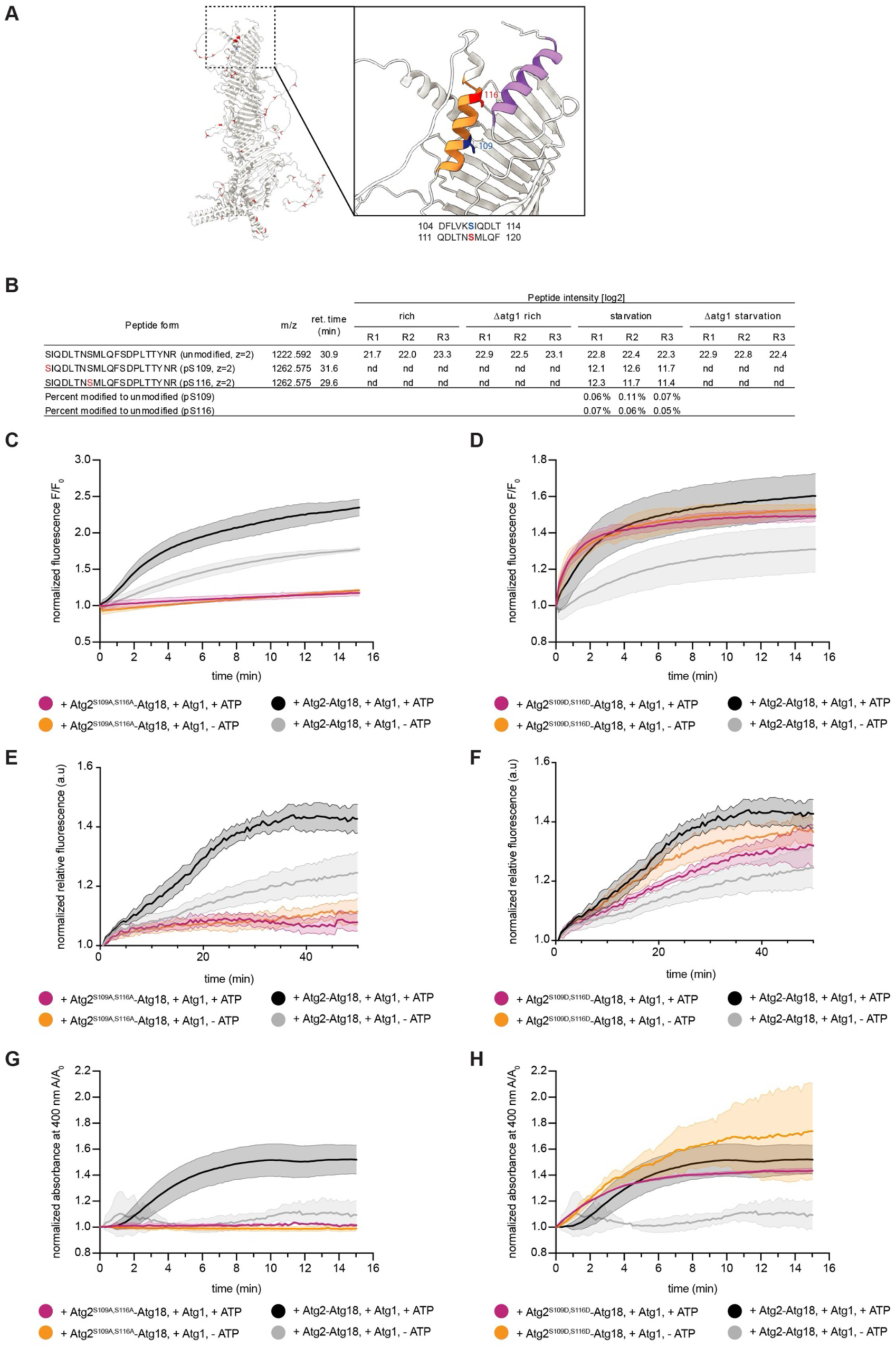
Phosphorylation of the Atg2 N-terminus enhances lipid transfer. (**A**) Atg2 AlphaFold model including Atg1-dependent phospho-sites (in red and blue) identified by mass-spectrometry. S109 (blue) and S116 (red) are located in helix H2 (orange) at the N-terminus within Atg1 kinase consensus motifs. Helix H1 is labelled in purple. (**B**) Relative peptide quantification across all conditions. The phosphor-peptides were only detected in wild type samples under starvation conditions. The percent ratio of the modified to the unmodified peptide suggests a low site occupancy but can be influenced by different ionization efficiencies or missed cleavage events during enzymatic digest. “nd”, not detected. (**C-D**) FRET-based lipid transfer assays of Atg2^S109A,S116A^-Atg18 (**C**) and Atg2^S109D,S116D^-Atg18 (**D**), showing lipid transfer activity compared to wild type Atg2-Atg18. Atg2^S109A,S116A^-Atg18 transfer activity is almost abolished while Atg2^S109D,S116D^-Atg18 showed constitutive activation. Data are means ± SD (n = 3). (**E-F**) Microscopy-based lipid transfer assay with GUVs and liposomes of Atg2^S109A,S116A^-Atg18 (**E**) and Atg2^S109D,S116D^-Atg18 (**F**). Data are means ± SEM (n ≥ 3). Wild type curves are the same as in Fig. 1I. (**G-H**) Absorbance-based turbidity assay to test membrane tethering activity of Atg2^S109A,S116A^-Atg18 (**G**) and Atg2^S109D,S116D^-Atg18 (**H**). Tethering was abolished with Atg2^S109A,S116A^ and comparable to activated wild type Atg2-Atg18 with Atg2^S109D, S116D^-Atg18. Data are means ± SD (n = 3).

**Table 1.** Generated mutations in Atg2^43xA^, Atg2^N-10xA^, and Atg2^C-12xA^.

| construct | Mutated sites |
|---|---|
| Atg2<sup>43xA</sup> | S95A, S96A, S98A, S103A, T114A, S116A, S138A, S140A, S142A, S146A, S249A, T258A, S259A, S263A, S269A, S307A, S624A, S629A, S635A, S636A, S762A, S769A, S791A, S940A, S954A, S960A, S974A, S977A, S981A, S986A, S1001A, S1071A, S1074A, S1079A, S1086A, S1113A, S1115A, S1118A, S1293A, S1357A, S1368A, S1379A, T1413A |
| Atg2<sup>N-10xA</sup> | S95A, S96A, S98A, S103A, T114A, S116A, S138A, S140A, S142A, S146A |
| Atg2<sup>C-12xA</sup> | S1071A, S1074A, S1079A, S1086A, S1113A, S1115A, S1118A, S1293A, S1357A, S1368A, S1379A, T1413A |

To test whether phosphorylation at S109 and S116 is sufficient for stimulating lipid transfer, we generated phosphomimetic mutants by substituting the residues with aspartic acid. The single mutants Atg2^S109D^ and Atg2^S116D^ showed increased lipid transfer compared to wild type already in the absence of active Atg1 (“-ATP”). However, transfer still reached higher levels in the presence of ATP (Fig. EV1H,I). When both serines were replaced by aspartic acid, the transfer reached similar levels as the stimulated wild type already in absence of ATP and could not be further stimulated (Fig. 2D). Similar results were also observed with GUVs as acceptors and liposomes as donors in the microscopy-based FRET-assay (Fig. 2E,F, Fig. EV1J).

Next, we tested the tethering activity of the mutants with the turbidity-based assay. The results correlated well with the transfer assays. Liposomes did not cluster in the presence of Atg2^S109A,S116A^ independent of its phosphorylation state (Fig. 2G). The phosphomimetic Atg2^S109D,S116D^-mutant boosted liposome tethering already in the absence of active Atg1 (“-ATP”) (Fig. 2H). Mutation of S109 to A or D correlated well with the respective double mutant in the tethering assay, which was not the case for the analogous S116 mutants (Fig. EV1K-N).

These results show that Atg1 stimulates lipid transfer by Atg2 by phosphorylating S109 and S116.

### Phosphorylation of the Atg2 N-terminus destabilizes helix H2 and promotes lipid access to the hydrophobic groove

To elucidate the molecular mechanism of Atg2 activation by phosphorylation, we combined structure prediction with multiscale molecular dynamics (MD) simulations.

We first predicted the structure of full-length Atg2 (UniProt P53855, residues 1–1592) with AlphaFold3 (Abramson *et al*, 2024), specifying S109 and/or S116 as phosphoserines in the prediction (three predictions with different seeds and five models each per variant). Despite a similar fold in all models, the placement of the N-terminal amphipathic helices H1 (residues 6–21) and H2 (residues 104–118) varied among the different variants. In predictions of Atg2^WT^ and Atg2^S116ph^, H1 and H2 occlude the N-terminus (Fig. EV2). Predicted models of Atg2^S109ph^ feature varying degrees of H2 displacement, resulting in a progressively more open hydrophobic cavity. Most models of Atg2^S109ph,S116ph^ show an extreme displacement of H2. It has to be noted, however, that the confidence of the predictions for H1 and H2 is relatively low, especially for the phosphorylated variants (Appendix Fig. S2A,B).

To assess the stability of H1 and H2 in the predicted models, we performed all-atom (AA) microsecond-long simulations of the N-terminal region (residues 1–230) in solution, starting from two conformations taken from the full-length AlphaFold3 models: the closed conformation of the rank-1 Atg2^WT^ model and the more open conformation of the rank-2 Atg2^S109ph^ model, in which H1 and H2 are slightly displaced from the groove. In the simulations, the closed structures were stable. By contrast, the open structures show a displacement of H2 towards the hydrophobic groove of all phosphorylation states, with H2 adopting a position and orientation that only partially overlaps with that of the closed structure (Appendix Fig. S2C). This displacement further suggests variability in the position of H2, consistent with the comparably low confidence scores in AlphaFold3 predictions.

To assess whether a rearrangement of H1 and H2 affects lipid access, we performed coarse-grained (CG) MD simulations of full-length Atg2 in solution with randomly dispersed POPC lipids, starting from five different AlphaFold3 models of each variant (ten 1-µs simulations per model, 50 per variant). Under these conditions, lipids rapidly self-assemble into dynamic micelles and bicelles, and also access the hydrophobic groove of Atg2. CG-MD simulations of Atg2^S109ph,S116ph^ showed the highest N-terminal lipid access, followed by Atg2^S109ph^. In Atg2^WT^ and Atg2^S116ph^, where the cavity is partially occluded by H1 and H2, lipid uptake from the N-terminus was rare (Fig. 3A).

**Figure 3.**
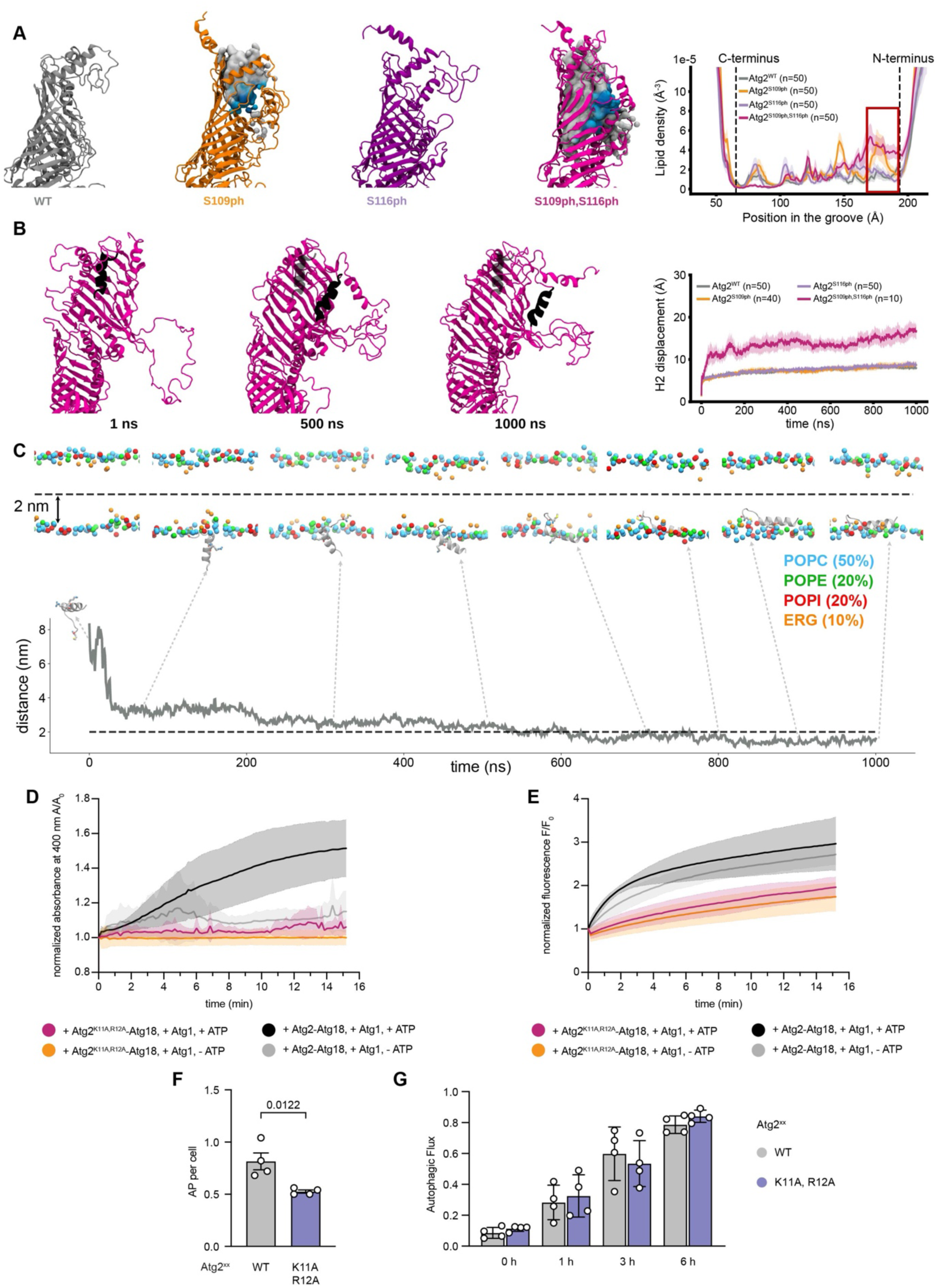
Phosphorylation of the Atg2 N-terminus destabilizes helix H2 and promotes lipid access to the hydrophobic groove in multiscale MD simulations. **(A)** Phosphorylation promotes lipid access to the N-terminal end of the hydrophobic groove in coarse-grained (CG) MD simulations of full-length Atg2 (residues 1–1592) with dispersed POPC lipids in solution. Left: N-terminal end of the groove in one representative simulation per variant (in which the density profile is closest to the mean), backmapped to atomistic resolution. Lipids that spent at least 10 ns in the N-terminal region of the groove are shown as surfaces (gray tails, blue head groups) at every 10th ns of their residence. Right: density of lipid head groups (PO4 beads) along the hydrophobic groove. Curves are means of 50 simulations per variant (ten 1-µs simulations for each of the five AlphaFold3 models), with shaded regions indicating the standard error of the mean (SEM). The density was calculated in a 30 Å × 30 Å × 220 Å box (1 Å bins along the groove) centered on the protein’s center of mass. Dashed lines mark the C- and N-terminal ends of the groove; the red box marks the N-terminal region (168–196 Å). (**B**) Displacement of H2 in CG simulations starting from AlphaFold models without H2 displacement. Left: snapshots of the Atg2^S109ph,S116ph^ rank-1 simulation with the largest H2 displacement at 1, 500 and 1000 ns (backmapped, superposed on the N-terminal core of the 1-ns frame). H2 is shown in black, its position at 1 ns is shown as a transparent helix. Right: displacement of H2 (residues 104–118) from its position in the starting AlphaFold3 model, after superposition of the N-terminal core (residues 35–100 and 176–229), as a function of time. Only simulations started from models with H2 in place are included (Atg2^WT^ and Atg2^S116ph^, ranks 1–5, n = 50 each; Atg2^S109ph^, ranks 2–5, n = 40; Atg2^S109ph,S116ph^, rank 1, n = 10). Data are means with shaded regions indicating SEM. (**C**) The H1 helix spontaneously inserts into the yeast ER membrane in AA simulations, as shown by the time evolution of the vertical distance (Z-axis) between the center of mass (COM) of H1 and the bilayer midplane. A distance below 2 nm (dashed line) is defined as membrane insertion. The trajectory represents a single simulation replicate, snapshots along this simulation are displayed (dashed arrows). Headgroup phosphates and the hydroxyl groups of ergosterol are shown as spheres. Positively charged residues in the protein (M1, K11 and R12) are highlighted as sticks. (**D**) Tethering was almost abolished with Atg2^K11A,R12A^ when compared to wild type Atg2-Atg18. Data are means ± SD (n = 3). (**E**) FRET-based lipid transfer assays of Atg2^K11A,R12A^-Atg18 showing that lipid transfer activity was reduced when compared to wild type Atg2-Atg18. Data are means ± SD (n = 3). (F) Ǫuantification of Atg8 positive autophagosome from live widefield microscopy of Atg2^WT^ and Atg2^K11A,R12A^ cells expressing 2xGFP-Atg8 during nitrogen starvation (1h). Data are means ± SEM (n = 4, 50 cells each), nested t-test analysis. (**G**) Autophagic flux of Atg2^WT^ and Atg2^K11A,R12A^ cells expressing 2xGFP-Atg8 after 0,1, 3, and 6 h of starvation. Data are means ± SD (n = 4).

To assess whether the increase in lipid uptake depends on phosphorylation or on H2 displacement, we separately analysed simulations starting from models where H2 was extremely displaced (one model of Atg2^S109ph^ and four models of Atg2^S109ph,s116ph^). This analysis showed a strong correlation between H2 displacement in the starting structure and N-terminal lipid uptake (Fig. EV2B). The CG-MD simulations thus show that a displaced H2 opens the N-terminal entrance of the hydrophobic groove for lipids.

To test whether phosphorylation can drive a displacement of H2, we followed the position of this helix in the CG-MD simulations starting from models with H2 close to its position in Atg2^WT^. In Atg2^S109ph,S116ph^, H2 moved away from its predicted position within the simulation time, whereas Atg2^S109ph^, Atg2^S116ph^, and Atg2^WT^ showed no substantial displacement (Fig. 3B, Movie EV1). This is consistent with a model in which double phosphorylation promotes the displacement of H2, increasing lipid uptake. Together, our simulations support a mechanism in which phosphorylation of S109 and S116 destabilizes H2 and promotes its displacement, resulting in an increase in lipid uptake. Given that H1 is positioned at the N-terminus and contains conserved basic residues (K11, R12) and considering the increase in membrane tethering by phosphorylated Atg2 (Fig. 1G), we hypothesized that it may play a dual role in both gating the hydrophobic groove and tethering Atg2 to membranes. To test this, we performed AA simulations of the N-terminal region of Atg2 (residues 1–20, including a short, disordered N-terminal segment and the H1 helix) in an ER-mimicking bilayer. In all three replicates, this fragment spontaneously inserted into the membrane. The N-terminus initiated contact with the lipid headgroups, followed by reorientation of the H1 helix, leading to stable membrane insertion (Fig. 3C).

We replaced the basic residues K11 and R12 with alanines and assessed membrane binding, lipid transfer and autophagy in vitro and in cells. The mutant Atg2^K11A,R12A^-Atg18 showed a strong defect in membrane binding as well as lipid transfer in the in vitro lipid transfer assay (Fig. 3D,E). In cells, the number of autophagosomes was reduced (Fig. 3F), but autophagic flux was not affected (Fig. 3G). However, consistent with a role of H1 in membrane binding, a recent study found an autophagic defect upon deletion of the first 12 amino acids (Kotani *et al*, 2026).

Together, our data suggests that H1 can insert into the donor membrane supporting a dual function of this helix as both gating element and membrane anchor.

### Phosphorylation of the Atg2 N-terminus promotes efficient phagophore formation

We proceeded to test how the phosphorylation of the Atg2 N-terminus affects autophagy. Accordingly, we generated strains expressing wild type Atg2 or the non-phosphorylatable S109A and S116A variants. Expression levels of the Atg2-variants were comparable to the wild type, and they also correctly localized to the tip of the phagophore in starved strains co-expressing BFP-prApe1 and 2xmCherry-Atg8 (Fig. EV3A,B). To examine the impact on phagophore expansion, we inducibly overexpressed BFP-prApe1 in 2xGFP-Atg8 cells during nitrogen starvation (Fig. 4A). PrApe1 overexpression leads to the generation of giant oligomers that are targeted but not cleared by autophagy, allowing the monitoring of phagophore expansion by live microscopy. We found that the maximal membrane expansion and the phagophore growth rate were significantly reduced in cells expressing Atg2^S109A^ compared to wild type. While Atg2^S116A^ showed a minor defect, the double mutant Atg2^S109A,S116A^ showed an even stronger expansion defect compared to Atg2^S109A^ (Fig. 4B,C). These results suggest that the phosphorylation of the Atg2 N-terminus at positions 109 and 116 is required for efficient phagophore expansion. To assess if this defect impacts autophagic flux, we monitored processing of 2xGFP-Atg8 via western blot analysis (Fig. 4D). Consistent with the observed phagophore expansion defect, expression of Atg2^S109A,S116A^ significantly impairs autophagic flux.

**Figure 4.**
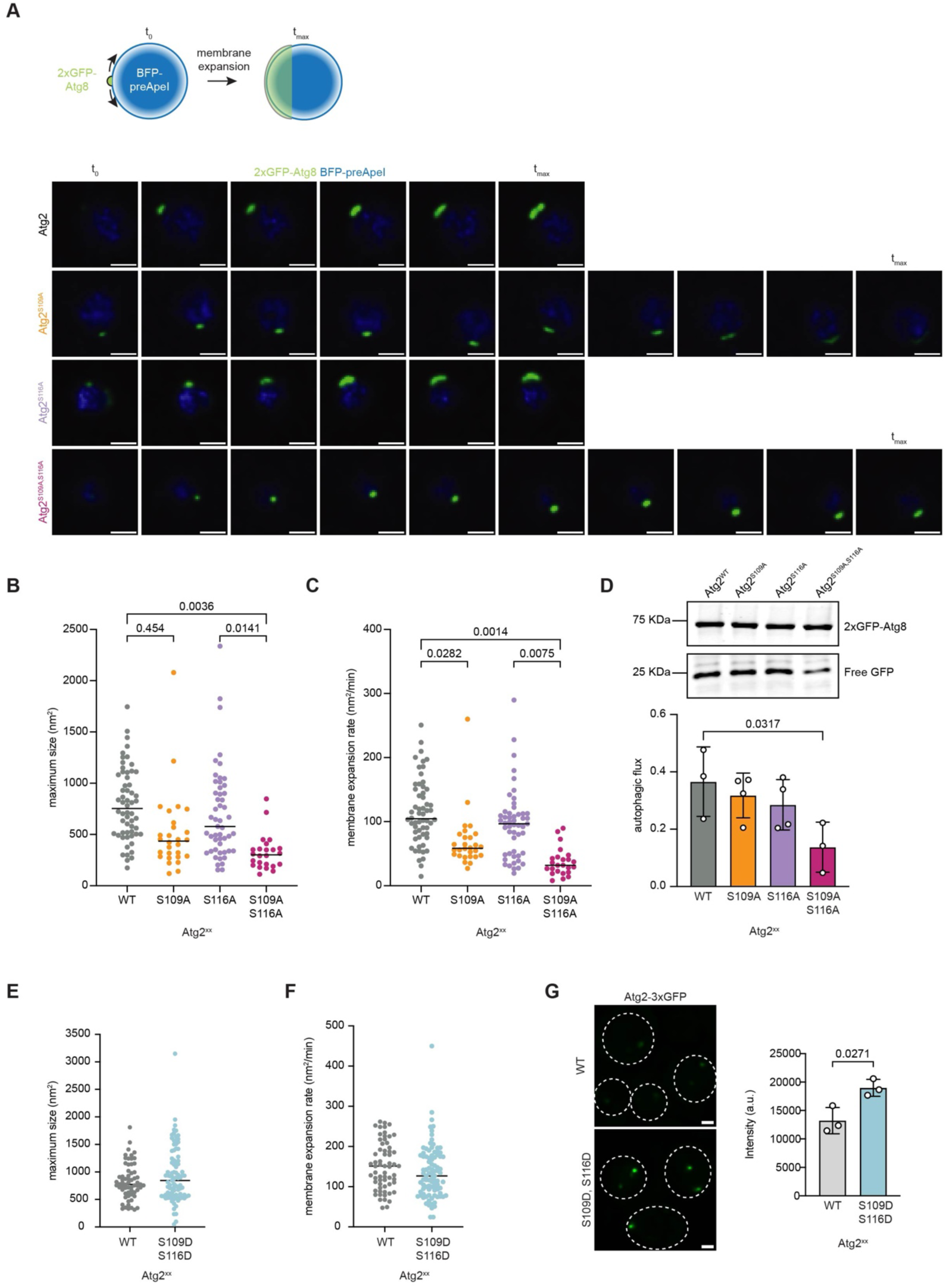
Impaired phosphorylation of the Atg2 N-terminus leads to inefficient autophagosome formation. **(A)** Schematic representation of the Giant ApeI assay. ApeI overexpression generates the formation of large oligomers subsequently targeted by autophagy. Measuring the size of the Atg8 positive membrane over time allows to calculate the expansion rate and maximal membrane expansion (top). Representative time course of indicated strains subjected to Giant ApeI assay (bottom). Scale bar: 1 µm. (**B-C**) Ǫuantification of maximal phagophore expansion (**B**) and membrane expansion rate (**C**) of indicated strains expressing 2xGFP-Atg8 and overexpressing the copper-inducible BFP-preApeI. Nested One-way ANOVA, n = 3 (median shown). (**D**) Autophagic flux of indicated strains expressing 2xGFP-Atg8 after 3 h of starvation. Flux was significantly reduced in Atg2^S109A,S116A^ cells. Data are means ± SD (n ≥ 3). One-way ANOVA. (**E-F**) Ǫuantification of maximal phagophore expansion (**E**) and membrane expansion rate (**F**) of indicated strains, as in (**B-C**) (n = 3, median shown). (**G**) Left, live widefield microscopy of endogenously 3xGFP-tagged Atg2 in cells expressing 2xmCherry-Atg8. Cells were imaged after shift to nitrogen starvation to induce autophagy (1 h). Pictures are maximal signal intensity Z projections. Scale bar: 1µm. Right, respective quantification of average Atg2 intensity. Data are means ± SD (n = 3, 50 structures per experiment), nested t-test.

We then proceeded testing the effects of expression of phosphomimetic Atg2^S109D,S116D^ during autophagy. Atg2^S109D,S116D^ did not affect maximal size nor autophagosome expansion rates in cells overexpressing BFP-prApe1 during nitrogen starvation (Fig. 4E,F), as well as autophagic flux (Fig. EV3C). However, fluorescent Atg2^S109D,S116D^ formed brighter structures compared to Atg2^WT^ during autophagosome biogenesis (Fig. 4G) suggesting increased recruitment or residence time at the PAS. Taken together, these results show that the phosphorylation of Atg2 at its N-terminus accelerates phagophore expansion and therefore promotes efficient autophagic flux.

### The VAP family protein Scs2 interacts with the Atg1 complex subunit Atg13

Lipid transfer via shuttles or BLTPs at MCSs is often supported by ER-embedded VAP family proteins (Reinisch *et al*, 2025; Prinz *et al*, 2020). In autophagy, they were shown to recruit Atg2 as well as the ULK1/Atg1 kinase complex to the ER (Du *et al*, 2026; Zhao *et al*, 2018; Kotani *et al*, 2026). To elucidate the role of the VAP family protein Scs2 on autophagosome biogenesis, we first monitored its localization during nitrogen starvation. Genomically mCherry-tagged Scs2 marked the ER and localized in close proximity to Atg8-positive structures (Fig. 5A). We excluded Scs2 as autophagy cargo via western blot analysis (Fig. EV4A). Next, we tested the effect of *SCS2* deletion on autophagy. After one hour of nitrogen starvation cells lacking Scs2 formed significantly fewer autophagosomes. At the same time, the number of Atg8-positive puncta tended to increase, suggesting impaired autophagosome biogenesis (Fig. 5B). Consistently, autophagic flux was reduced in *Δscs2* cells (Fig. 5C). To exclude contribution of the Scs2 paralogue Scs22, we performed the same experiments in *Δscs22* and *Δscs2Δscs22* cells. These experiments indicate that only Scs2 and not Scs22 is involved in autophagosome biogenesis (Fig. EV4B,C).

**Figure 5.**
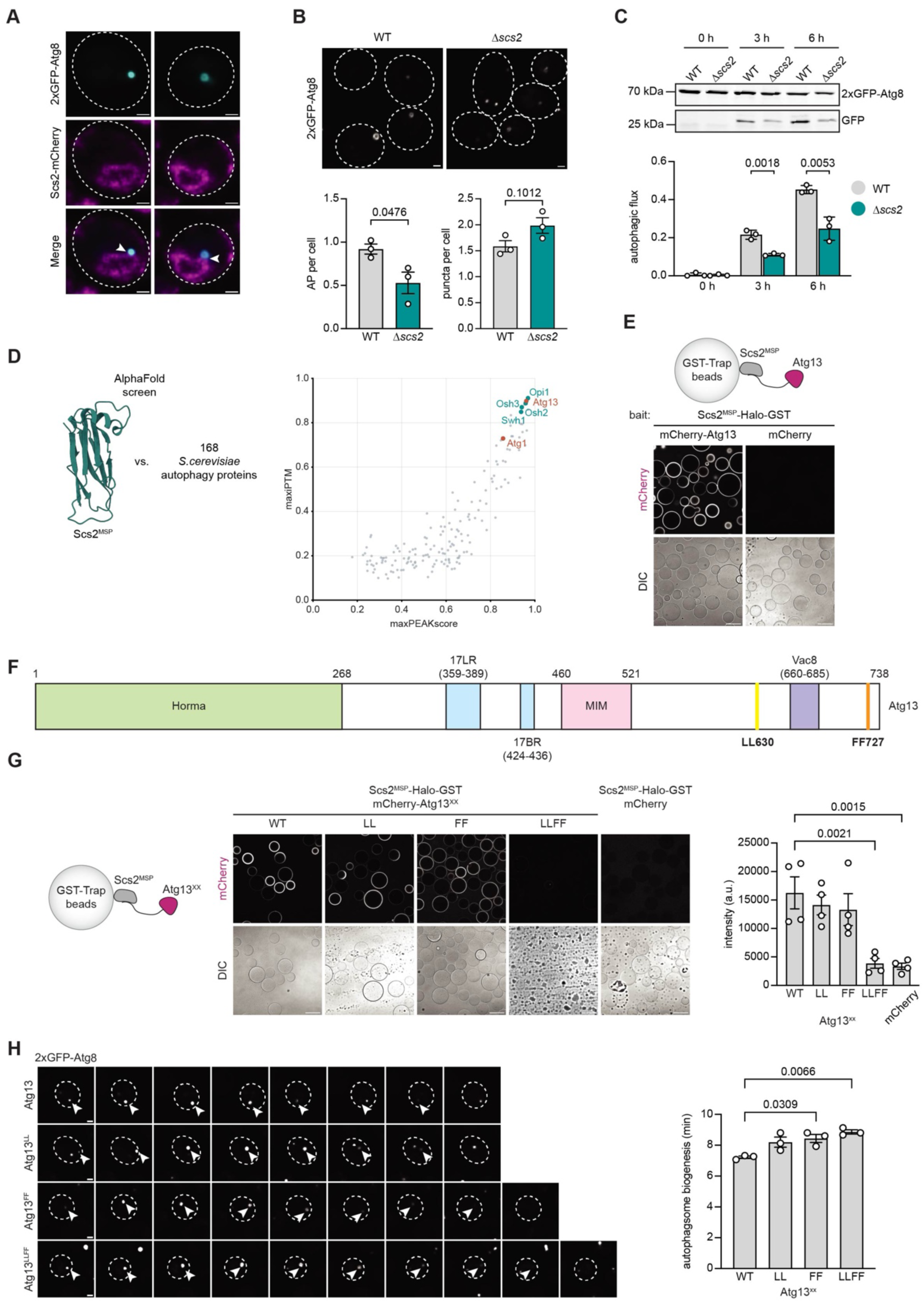
The VAP family protein Scs2 interacts with the Atg1 complex subunit Atg13. **(A**) Live widefield microscopy of endogenously mCherry-tagged Scs2 in cells expressing 2xGFP-Atg8. Cells were imaged after shift to nitrogen starvation to induce autophagy (1 h). Ring structures positive for Atg8 are classified as autophagosomes while non-ring structures are immature autophagosomes/phagophores. Scale bar: 1µm. (**B**) Live widefield microscopy of wild type and *Δscs2* cells expressing 2xGFP-Atg8 during nitrogen starvation (55 min, top) and quantification (bottom) of Atg8 positive autophagosome (left) and puncta (right) in wild type, and *Δscs2* cells. Data are means ± SEM (n = 3, 50 cells each), nested t-test analysis. (**C**) Autophagic flux of indicated strains expressing 2xGFP-Atg8 after 0, 3, and 6 h of starvation. Data are means ± SD (n = 3). Unpaired t-test. (**D**) Structural representation of the Scs2 MSP domain used for the AlphaFold based *in silico* pull-down (left) and graphical representation of the *in silico* pull-down results (right). Positive controls in cyan, selected preys in red. (**E**) Microscopy-based pull-down assay where purified Scs2^MSP^-Halo-GST was immobilized on beads as bait and incubated with mCherry-Atg13 (n = 3). (**F**) Schematic representation of Atg13 protein and its domains (Fujioka *et al*, 2014; Yamamoto *et al*, 2016; Jao *et al*, 2013; Park *et al*, 2020; Scott *et al*, 2000). Amino acid residues contained in FFAT motifs responsible for binding to Scs2 are highlighted in yellow and orange. (**G**) Microscopy-based pull-down assay where purified Scs2^MSP^-Halo-GST was immobilized on beads as bait and incubated with mCherry-Atg13 mutants or mCherry as control. Data are means ± SEM, nested One-way ANOVA, n = 4. (**H**) Representative time course of indicated 2xGFP-ATG8 strains expressing Atg13 variants (left) and respective quantification of autophagosome biogenesis duration (right). Arrowheads indicate nascent autophagosomes. Scale bar: 1 µm. Data are means ± SEM, nested One-way ANOVA vs wild type, n = 3 (15 events per strain each). Scale bars of microscopy-based pull-down assays: 100 µm.

Next, we asked if Scs2 could recruit components of the upstream autophagy factors, including the Atg1 kinase complex. We therefore performed an AlphaFold2 multimer screen with the cytosolic major sperm protein (MSP) domain of Scs2 and 168 autophagy-associated proteins. Known interactors served as positive controls. Among the top hits was the Atg1 complex subunit Atg13. We also found Atg1 in the AlphaFold2 multimer screen, albeit with lower confidence (Fig. 5D).

To test the interaction between Scs2 and Atg13 *in vitro* we purified the MSP domain of Scs2 and bound it to beads to image fluorescently labeled Atg13 recruitment by microscopy. Confirming the AlphaFold screen, we observed recruitment of Atg13 to the Scs2^MSP^-coated beads (Fig. 5E, Fig. EV4D).

We conclude that Scs2 binds Atg13 and has the potential to localize the Atg1 kinase in proximity to the ER.

### Atg13 interacts with Scs2 via a FFAT motif

Next, we set out to identify the Scs2 binding site(s) in Atg13. The MSP domain of VAP proteins binds interactors via an FFAT motif (two phenylalanines in an acidic tract) (Loewen *et al*, 2003). Indeed, AlphaFold3 predictions of Scs2 and Atg13 showed an interaction between Scs2 and the C-terminus of Atg13 involving F727 and F728 (Fig. 5F, Fig. EV4E (left)). To test whether this FFAT motif is responsible for the interaction, we generated an Atg13-mutant with F727 and F728 mutated to alanines (Atg13^FF^). When we repeated the microscopy-based pull-down with this mutant, however, we could not observe a reduced recruitment (Fig. 5G). AlphaFold3 predictions of Scs2 and Atg13^FF^ suggested a second interaction site between Scs2 and Atg13 further upstream with an acidic motif containing two leucines at positions 630 and 631 (Fig. 5F, Fig. EV4E (right)). Additional mutation of these two leucines (Atg13^FFLL^) abolished the interaction with Scs2^MSP^ *in vitro*, whereas replacing only L630 and L631 with alanines (Atg13^LL^) did not affect the recruitment to Scs2^MSP^ (Fig. 5G). We then proceeded to evaluate the Scs2-Atg13 binding mutants *in vivo*. To this end, we genomically integrated Atg13 variants in 2xGFP-Atg8 cells to follow autophagosome biogenesis. In line with our *in vitro* data, Atg13^LL^ did not affect autophagosome formation. Interestingly though, we detected about twice as many autophagosomes in cells expressing Atg13^FF^ or Atg13^FFLL^ (Fig. EV4F). Despite the elevated number of autophagosomes, autophagic flux was not significantly increased (Fig. 4G). This effect may be a result of slower autophagosome biogenesis (Dabrowski *et al*, 2023). We therefore monitored autophagosome biogenesis by time-lapse fluorescence microscopy. Indeed, autophagosome formation took longer in cells harboring Atg13^FF^ or Atg13^LLFF^ compared to wild type (Fig. 5H). Mutating L630 and L631 in combination with F727 and F728 did not result in an increased effect on autophagosome formation indicating that mutation of F272 and F728 are the primary cause of the defect *in vivo.* These findings show that Atg13 interacts with Scs2 via FFAT motifs in its C-terminus and that a loss of this interaction slows down autophagosome formation.

### Atg1 is recruited to Scs2 in a phosphorylation-dependent manner

Atg13 forms a complex with the Atg1 kinase (Stjepanovic *et al*, 2014; Suzuki *et al*, 2007). We therefore asked if Atg1 is recruited to Scs2 via Atg13 enabling it to phosphorylate autophagy proteins localizing at the ER including Atg2. We performed the Scs2^MSP^ – mCherry-Atg13 microscopy-based beads assay as above (Fig. 5E) and added GFP-Atg1. GFP-Atg1 was recruited to the Scs2^MSP^-coated beads. When we added ATP to enable Atg1 kinase activity, the recruitment of GFP-Atg1 to the Scs2^MSP^-coated beads was increased significantly (Fig. 6A). At the same time, the interaction between Scs2^MSP^ and mCherry-Atg13 was reduced (Fig. 6A), already at relatively low concentrations of GFP-Atg1 (Fig. EV5A-C). To exclude an effect on Atg1 binding from residual Atg13 bound to Scs2, we performed the experiment in the absence of Atg13. Notably, in the absence of Atg13, Atg1 bound to Scs2^MSP^ only in the presence of ATP and the kinase-deficient Atg1^D211A^ did not interact with Scs2^MSP^ (Fig. 6B). Taken together, this suggests competitive binding of Atg1 and Atg13 to Scs2 with a dominance of Atg1 over Atg13. In addition, there could be other changes in Atg13 with regard to Scs2 binding elicited by phosphorylation. The lack of indirect binding of Atg13 to Scs2 via Atg1 is likely due to the weakening of the Atg1 – Atg13 interaction by Atg1 activity (Schreiber *et al*, 2021).

**Figure 6.**
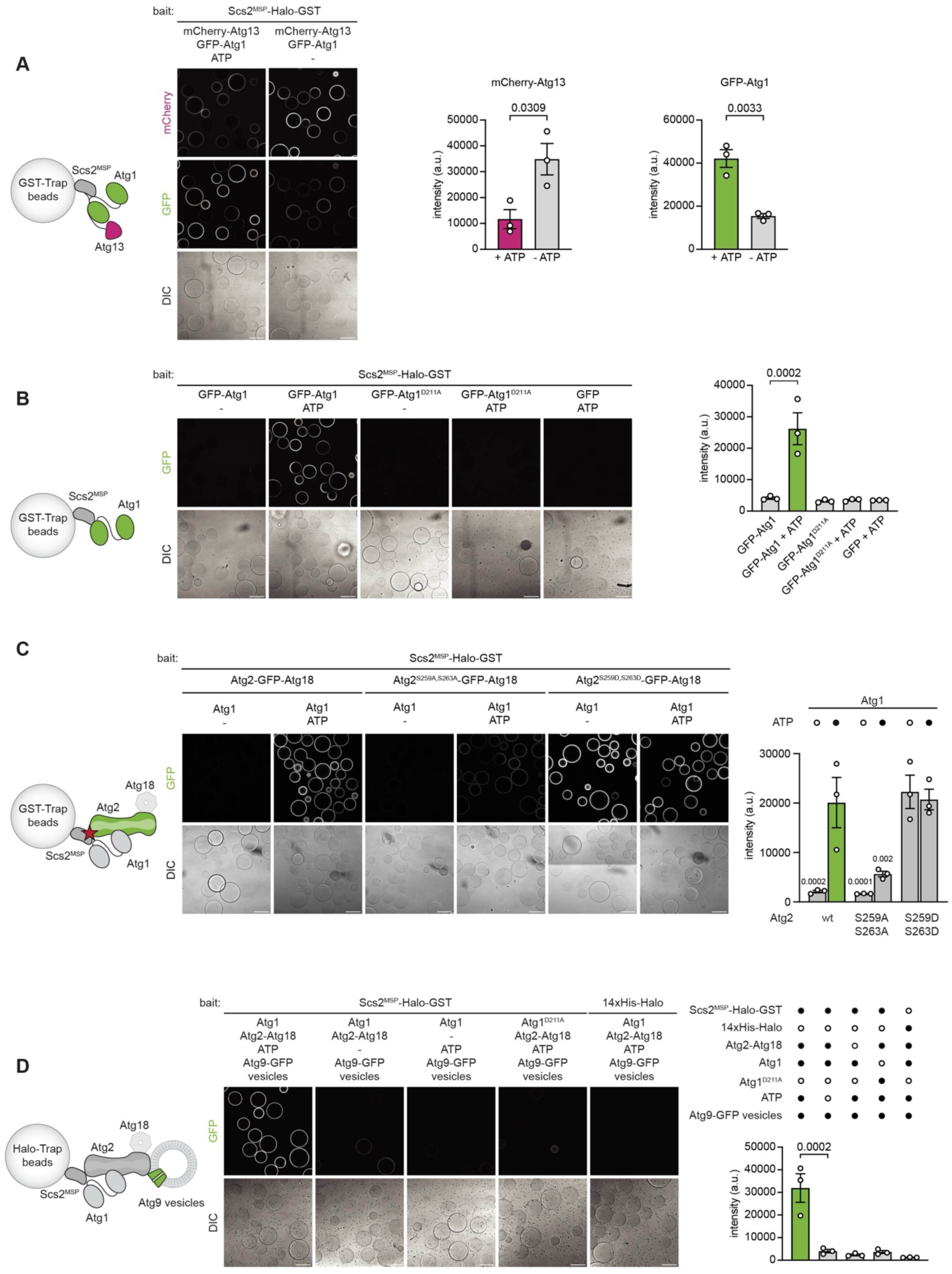
Reconstitution of a minimal MCS including Scs2, Atg1, Atg2, and AtgG vesicles. (**A**) Microscopy-based pull-down assay where purified Scs2^MSP^-Halo-GST was immobilized on beads as bait and incubated with mCherry-Atg13 and GFP-Atg1 in presence or absence of ATP. Data are means ± SEM (n = 3), nested t-test. Fig. 6A and Fig. EV5B were generated analyzing the same dataset and representative pictures are partially repeated. (**B**) Microscopy-based pull-down assay where purified Scs2^MSP^-Halo-GST was immobilized on beads as bait and incubated with GFP-Atg1, GFP-Atg1^D211A^ or GFP as control in presence or absence of ATP. Data are means ± SEM (n = 3), nested One-way ANOVA vs GFP-Atg1. (**C**) Microscopy-based pull-down assay where purified Scs2^MSP^-Halo-GST was immobilized on beads as bait and incubated with wild type Atg2-GFP, Atg2^S259A,S263A^-GFP or Atg2^S259D,S263D^-GFP, Atg1, and in presence or absence ATP. Data are means ± SEM (n = 3), nested One-way ANOVA vs Atg2-GFP—Atg18 + Atg1 + ATP. (**D**) Microscopy-based pull-down assay where purified Scs2^MSP^-Halo-GST was immobilized on beads as bait and incubated with Atg9-GFP vesicles in presence or absence of Atg2, Atg1, and ATP. 14xHis-Halo was used as control. Data are means ± SEM (n= 3), nested One-way ANOVA vs Atg1 Atg2—Atg18 Atg9-EGFP. Scale bars, 100 µm.

### Atg1 dependent contact site formation with AtgG vesicles

From our data, we conclude that the BLTP Atg2 is a substrate for ER-associated Atg1. We have shown that the membrane association of Atg2 is enhanced upon phosphorylation by Atg1 (Fig. 1G), and that Atg1 interacts with Scs2 (Fig. 6B). Other BLTPs were previously shown to interact with VAP family proteins (Du *et al*, 2026; Kumar *et al*, 2018; Guillén-Samander *et al*, 2021; Kotani *et al*, 2026). Recently Atg2 also was shown to bind to Scs2 via a phospho-FFAT motif *in vivo* (Kotani *et al*, 2026). We therefore set out to test this interaction in a minimal system *in vitro*. We coated beads with Scs2^MSP^ and monitored recruitment of Atg2-GFP by fluorescence microscopy. In accordance with Kotani et al. (Kotani *et al*, 2026), Atg2-GFP bound Scs2 only in the presence of Atg1 and ATP, demonstrating a phosphorylation-dependent activation of this interaction (Fig. EV5D). Mutation of S259 and S263 in the reported phospho-FFAT motif to alanines resulted in the loss of interaction independent of Atg1 activity (“+ ATP”) whereas the phosphomimetic mutant Atg2^S259D,S263D^ was recruited to the Scs2^MSP^-coated beads already in the absence of ATP or Atg1 (Fig. 6C, Fig. EV5E). We assessed the effect of the phosphorylation on S259 and S263 in cells by mutating these sites to alanines. Cells expressing Atg2^S259A,S263A^ showed a reduction of autophagosomes but no autophagic flux defect (Fig. EV5F,G). Phagophore expansion was monitored by the giant ApeI assay (Fig. 4A) and revealed no effect on the size of phagophores but decreased expansions rates (Fig. EV5H,I) in contrast to cells expressing Atg2^S109A,S116A^ that displayed a strong impairment in autophagic flux, phagophore expansion and expansion speed (Fig. 4B-D). Finally, we set out to reconstitute a minimal contact site and tested whether Atg9 vesicles, the seed membranes for the expanding phagophore at ER-autophagosome contact sites (Sawa-Makarska *et al*, 2020; Olivas *et al*, 2023), could be recruited via the Scs2-Atg2 axis. Indeed, Atg9-GFP vesicles purified from cells localized to Scs2^MSP^-coated beads upon addition of Atg2, Atg1 and ATP (Fig. 6D). Thus, establishing a contact between Scs2 and Atg2 is sufficient to tether Atg9 vesicles to the ER.

Collectively, our results show that the Atg1 kinase can promote the membrane contact site formation between the ER and Atg9 vesicles during autophagy initiation. They further show, how its kinase activity can simultaneously facilitate membrane binding and lipid transfer by the BLTP Atg2.

## Discussion

How BLTPs are spatiotemporally regulated at MCSs is still incompletely understood (Reinisch *et al*, 2025). This is an important question because once BLTPs connect two membranes, they can rapidly equilibrate the lipids between them. It is therefore imperative to recruit them swiftly when needed, such as during autophagy initiation or membrane damage (Wang *et al*, 2025; Tan & Finkel, 2022; Cross *et al*, 2023). At the same time, spurious activation must be prevented.

Here, we show that the phosphorylation of a BLTP regulates its lipid binding and transfer activity. The phosphorylation of Atg2 is accomplished by the autophagy master regulator kinase Atg1. Previously it was shown that Atg1 phosphorylates Atg2 during autophagy but the role of this modification remained unresolved (Papinski *et al*, 2014; Hu *et al*, 2019). We found that the phosphorylation of S109 and S116 residing in helix H2 at the N-terminus of Atg2 leads to increased membrane tethering. Concomitant lipid transfer activity is increased by the opening of the entry to the N-terminal, ER-associated end of the lipid transfer groove. Interestingly, when the cryo-EM structure of the *Schizosaccharomyces pombe* Atg2 N-terminus was solved, it was observed that the helix H4, which corresponds to H2 in *Saccharomyces cerevisiae*, blocks the hydrophobic groove preventing lipid transfer (Osawa *et al*, 2019). Indeed, our MD simulations suggest a displacement of the helices H1 and H2 upon phosphorylation of S109 and S116 allowing lipid uptake. The introduction of two negatively charged phosphate groups likely drive electrostatic repulsion within the N-terminus, thereby promoting a conformational opening of the groove. We recently proposed a similar destabilization of the N-terminal helices to facilitate membrane binding and accelerate lipid transfer for the mammalian homologue ATG2A (preprint: Cabezudo *et al*, 2025).

Notably, Atg2 retains partial lipid transfer activity in the absence of Atg1-mediated phosphorylation *in vitro*, suggesting that the protein dynamically switches between open and closed conformations. We propose that phosphorylation of S109 and S116 shifts this equilibrium toward the open, active state, thereby stimulating lipid transfer activity. The local concentration of Atg1, the interaction with Scs2 and membrane insertion of H1 may render the open conformation long lived in cells.

Our data also seem to indicate a hierarchical role of S109 over S116 as Atg2^S116A^ retains a higher tethering capacity than Atg2^S109A^ *in vitro* (Fig. EV1K,L). In line with this idea, Atg2^S109A^ alone impairs phagophore expansion *in vivo* (Fig. 4B,C).

Our model is supported by parallel findings by Sakai and colleagues who have independently identified residues S109 and S116 as targets of Atg1-mediated activation of membrane binding, lipid transfer and phagophore biogenesis (preprint: Sakai *et al*, 2025). Interestingly, cells expressing the Atg2^S109D,S116D^ mutant do not show an increased autophagic flux, suggesting that other activities such as lipid supply or scrambling may be rate limiting under these conditions.

In the present study we also investigated the initiation of the MCS between the ER and the phagophore and reconstituted a minimal contact site involved in autophagy. We found that the Atg1 kinase complex and the BLTP Atg2 bind the VAP family protein Scs2. VAP family proteins are associated with MCSs and were also shown to be involved in mammalian and yeast autophagy (Du *et al*, 2026; Zhao *et al*, 2018; Kotani *et al*, 2026). We propose a mechanism where the Atg1 kinase complex localizes to the ER at least in part by the interaction between the C-terminal FFAT motifs of Atg13 and Scs2 (Fig. 7). Local accumulation of Atg1 complex condensates formed by Atg13 – Atg17 interactions (Fujioka *et al*, 2014) at the ER activates the kinase facilitating the binding between Scs2 and Atg1 partially outcompeting Atg13-binding. Atg13 recruits Atg9 vesicles to these condensates to cluster them close to the ER (Suzuki *et al*, 2015). Upon activation of Atg1, Atg13 is phosphorylated, disrupting the connection between Atg1 and Atg13 (Schreiber *et al*, 2021). Hence, the binding of Atg1 to Scs2 maintains its location at the ER to coordinate the phosphorylation of Atg2 promoting binding to Scs2 (Kotani *et al*, 2026), membrane binding and lipid transfer (Fig. 7). The Atg1-mediated activation of Scs2 binding by Atg2 via the FFAT motif has been independently found by Kotani et al. (Kotani *et al*, 2026). Other factors such as the TRAPPIII complex may act alongside Atg2 in connecting the ER and the phagophore (Gómez-Sánchez *et al*, 2025). The localization of the substrate Atg2 in close proximity to Atg1 may be assisted by the interaction with Atg29, also a subunit of the Atg1 complex, as reported recently (Yasuda *et al*, 2026).

**Figure 7.**
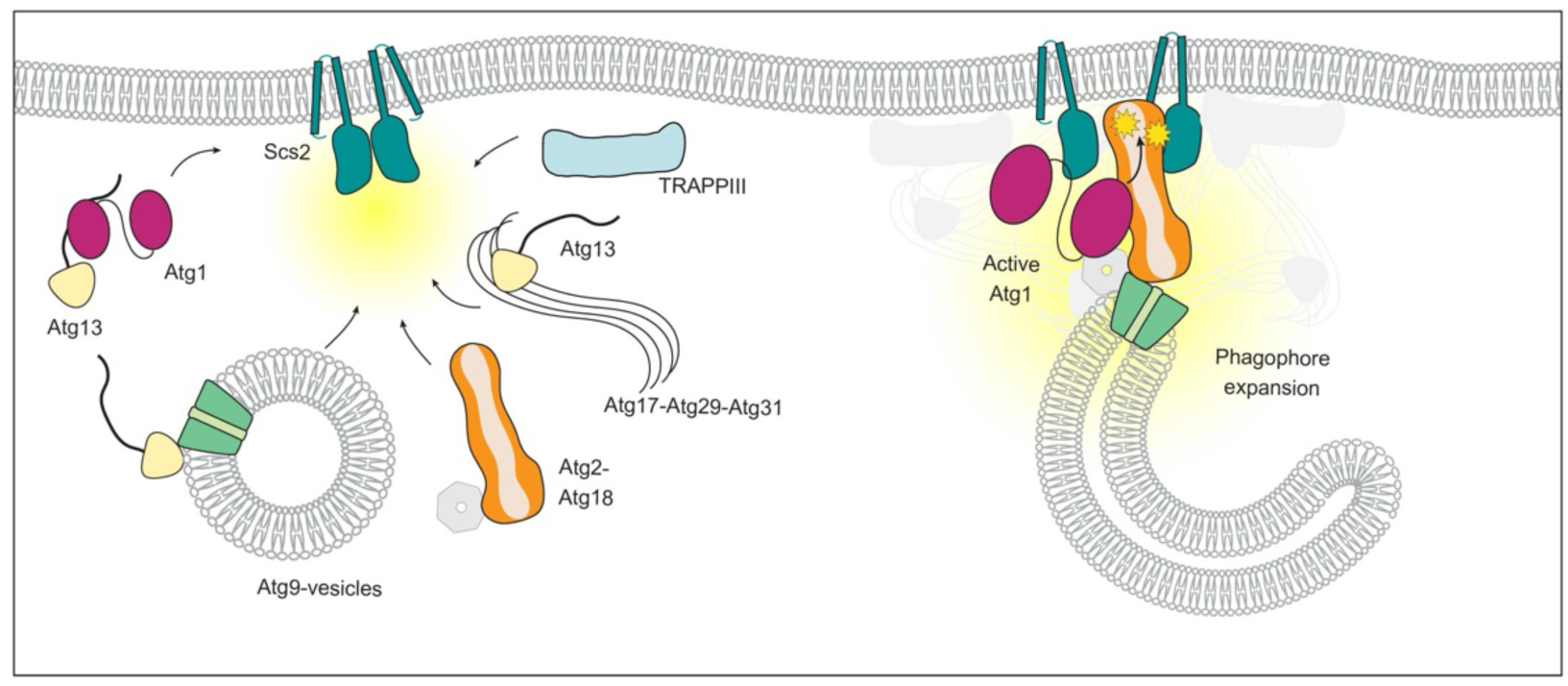
Proposed model for initiation of autophagosome biogenesis in vicinity to the ER. Local accumulation of Atg1 complex condensates formed by Atg13 – Atg17 interactions at the ER activates the Atg1 kinase facilitating the binding between Scs2 and Atg1 partially outcompeting Atg13 from Scs2. Atg13 recruits Atg9 vesicles to these condensates to cluster them close to the ER. Upon activation of Atg1, Atg13 is phosphorylated, disrupting the connection between Atg1 and Atg13. Hence, the binding of Atg1 to Scs2 maintains its location at the ER to coordinate the phosphorylation of Atg2 promoting its binding to Scs2, membrane binding and lipid transfer. Other factors such as the TRAPPIII complex may act alongside Atg2 in connecting the ER and the phagophore.

Our study reveals a new regulatory mechanism for the BLTP Atg2 that enhances lipid transfer at the ER–phagophore MCS. The Atg1-dependent phosphorylation of Atg2 couples kinase activation to lipid flux at the ER–phagophore contact site. Our study points toward a general mechanism for kinase-mediated regulation of BLTP activity at MCSs.

## Material and Methods

### Protein expression and purification

Atg2—Atg18 and Atg2-GFP—Atg18 variants were purified from genetically modified *S. cerevisiae* BY4741 strains listed in Table EV1. Cells were grown at 30°C in YP supplemented with 2% galactose to an OD600 between 6-10. After pelleting, cells were washed with cold H_2_O, with lysis buffer (200 mM NaCl, 50 mM Hepes pH 7.5, 1.5 mM MgCl_2_) and finally resuspended in lysis buffer containing cOmplete protease inhibitors (5056489001; Roche), an FY-inhibitor mix (39104; Serva), DNAse I (DN25; Sigma-Aldrich), benzonase, and 1 mM DTT. Resuspended cells were frozen in liquid nitrogen as pearls and lysed by freezer milling. The powder was thawed in lysis buffer containing cOmplete protease inhibitors, the FY-inhibitor mix and 1 mM DTT. Lysates were cleared by centrifugation (38,900 × *g* for 45 min at 4°C in a Hitachi Himac 20A rotor). The supernatant was incubated with IgG Sepharose 6 Fast Flow beads (17096901; Cytiva) on a rotary wheel for 1 h at 4°C. Beads with bound protein were washed twice with lysis buffer containing cOmplete protease inhibitors, the FY-inhibitor mix and 1 mM DTT, once with high salt buffer (700 mM NaCl, 25mM Hepes) and twice with lysis buffer. Bound protein was cleaved using TEV protease at 4°C overnight. The cleaved protein was concentrated, applied onto a Superose 6 Increase 10/300 size exclusion chromatography column (29091596; Cytiva) and eluted with elution buffer containing 150 mM NaCl, 50 mM Hepes pH 7.5, and 1 mM DTT. Fractions containing the purified proteins were pooled, concentrated, frozen in liquid nitrogen, and stored at -70°C.

For Fig. 1D the purification was split into two parts after the cleavage of the protein from the beads by TEV protease. The first part was incubated with 300 nM Atg1, 1 mM ATP and 1 mM MgCl_2_ while the second part was only incubated with Atg1 and MgCl_2_. Then both samples were applied to a Superose 6 Increase 10/300 size exclusion chromatography column (29091596; Cytiva) to separate Atg2-Atg18 from Atg1 and the ATP and eluted with elution buffer.

Atg1, Atg1^D211A^, GFP-Atg1, Atg1^1-454^, Atg13, mCherry-Atg13, mCherry-Atg13^FF^, mCherry-Atg13^LL^, mCherry-Atg13^LLFF^, and Atg2 were expressed in baculovirus-infected Sf9 insect cells via SMC1328, SMC1914, SMC1418, SMC2642, SMC1330, SMC1796, SMC2908, SMC3141, SMC3142, or SMC1908, respectively (Table EV2). Cells were harvested at 800 × *g* for 15 min, washed with PBS and flash frozen. For protein purification cells were lysed with a cell homogenizer in buffer (300 mM NaCl, 50 mM Tris pH 7.4 or 50 mM Hepes pH 7.5, 2 mM MgCl_2_) supplemented with 1 mM β-mercaptoethanol (8.05740; Sigma-Aldrich), benzonase, DNase I (DN25; Sigma-Aldrich), protease inhibitor cocktail (P8849; Sigma-Aldrich) and cOmplete protease inhibitor (5056489001; Roche). The lysate was cleared by centrifugation at 38,900 × *g* for 45 min (Hitachi Himac 20A rotor), and the supernatant was incubated with IgG Sepharose 6 Fast Flow beads (17096901; Cytiva) for 1 h at 4°C. The beads were then washed three times with buffer. Bound protein was eluted from the beads by TEV protease overnight at 4°C with buffer (300 mM NaCl, 50 mM Hepes pH 7.5, 1 mM DTT). The Atg2 construct was further processed by incubating the supernatant with glutathione sepharose beads (17075605; Cytiva) for 1 h at 4°C to remove the TEV protease. The supernatant was concentrated, flash frozen and stored at -70°C.

Atg1 and Atg13 constructs were concentrated and subjected to size exclusion chromatography on a Superdex 200 10/300 column (289990944; Cytiva). The protein was eluted in 150 mM NaCl, 25 mM Tris pH 7.4, 1 mM DTT. Peak fractions were collected, concentrated, flash frozen and stored at -70°C.

Scs2^MSP^-Halo-TEV-GST was purified from *Escherichia coli* Rosetta pLysS via SMC2728 (Table EV2). Cells were grown in lysogeny broth medium at 37°C until an OD600 of 0.4. Then expression was induced by the addition of 0.5 mM IPTG and further grown at 18°C overnight. The cells were harvested, resuspended in lysis buffer (300 mM NaCl, 50 mM Hepes pH 7.5, 2 mM MgCl_2_, 1 mM DTT, 2 mM β-mercaptoethanol (8.05740; Sigma-Aldrich), cOmplete protease inhibitors (5056489001; Roche), DNase I (DN25; Sigma-Aldrich)) and lysed by sonication. The lysate was cleared by centrifugation at 38,900 × *g* for 45 min at 4°C in a Hitachi Himac 20A rotor and the supernatant was incubated with glutathione sepharose beads (17075605; Cytiva) for 3 h at 4°C. The beads were washed and the protein was eluted in elution buffer (300 mM NaCl, 50 mM Hepes pH 7.5, 1 mM DTT, 40 mM reduced L-glutathione) overnight at 4°C. Eluted protein was concentrated and subjected to a Superdex S200 Increase 10/300 GL column (289990944; Cytiva) and eluted with elution buffer (150 mM Nacl, 25 mM Hepes pH 7.5, 1mM MgCl_2_, 1 mM DTT). Fractions containing the protein were pooled, concentrated, flash frozen and stored at - 70°C.

### Liposome preparation

Liposomes were prepared as described previously (Sawa-Makarska *et al*, 2020). In brief, lipids were mixed and dried under an argon stream. Donor liposomes for the microscopy-based lipid transfer assay in Fig. 1I and 2E-F contained 66% DOPC, 20% DOPE, 10% DOPS, 2% ATTO 425 DOPE (AD-425-161; ATTO-TEC), and 2% ATTO 540Ǫ DOPE (AD-540Ǫ-161, ATTO-TEC). Donor liposomes for the tethering assay in Fig. 2G,H and Fig. EV1K-N contained 66% DOPC, 24% DOPE, and 10% DOPS. All other donor liposomes contained 66% DOPC, 20% DOPE, 10% DOPS, 2% NBD-DPPE (810144C; Avanti Polar Lipids Inc.), and 2% lissamine rhodamine-DHPE (L-1392; Invitrogen) and acceptor liposomes contained 44% POPC, 6% POPS, 6% POPE, 41.5% liver PI, 2.5% PI3P. All percentages are in a w/v ratio. The lipids were dried under vacuum for 1 h. The lipid mixes were rehydrated with buffer (150 mM NaCl, 25 mM Tris pH 7.4) to reach a concentration of 1 or 2 mM and further sonication for 2 min in a water bath. The formed liposomes were extruded with a Mini Extruder (Avanti Polar Lipids Inc.) first through a 0.4 µm filter and then a 0.1 µm filter (10419504, 10417104; Whatman).

### GUV formation

Electroformation of GUVs was performed as described previously (Romanov *et al*, 2012). In brief, lipids (75% DOPC, 10% DOPS, 10% DOPE, and 5% PI3P (w/v)) were mixed at a concentration of 10 mg/ml in chloroform/methanol (3:1). The mixture was applied dropwise onto an indium-tin-oxide coated glass slide and further dried under vacuum for 3 h. The assembled electroformation chamber was filled with 300 mM sucrose and electroformation was performed at 30°C in three phases: Phase 1: 30 min, sine wave frequency f = 10 Hz, linear peak-to-peak amplitude ramp from 0.05 V to 1.41 V. Phase 2: 120 min, sine wave frequency f=10 Hz, peak-to-peak amplitude at 1.41 V. Phase 3: 30 min, square wave frequency f=4.5 Hz, peak-to-peak amplitude at 2.12 V. GUVs were diluted in GUV buffer (150 mM NaCl, 25 mM Tris pH7.4).

### FRET-based lipid transfer assays

The reaction contained 40 nM donor liposomes, 40 nM acceptor liposomes, 300 nM of Atg1 or Atg1 variants, 300 mM Atg13 where indicated, 1 mM ATP and 1 mM MgCl_2,_ in buffer (150 mM NaCl and 25mM Tris pH 7.4). The reaction was started with the addition of 400 nM (800 nM for Fig. 1C,D, 50 nM for Fig. 2D and Fig. EV1B and Fig. EVH,I) Atg2-Atg18 variants or 2 µM Atg2 (Fig. EV1A). Fluorescence intensity with an excitation wavelength of 485 nm and an emission wavelength of 535 nm was measured with a Tecan SPARK Multimode Microplate Reader (Tecan Life Sciences). For analysis, the individual measurements were divided by the starting value.

### Tethering assays

For the tethering assay 40 nM donors were mixed with 40 nM acceptors. The reaction contained 300 nM of Atg1, 1 mM ATP and 2 mM MgCl_2,_ in buffer (150 mM NaCl and 25mM Tris pH 7.4). The reaction was started with the addition of 800 nM (400 nM for Fig. 3D) Atg2-Atg18 variants. Absorption at 400 nm was measured with a Tecan SPARK Multimode Microplate Reader (Tecan Life Sciences). For analysis the individual measurements were divided by the starting value.

### *In vitro* kinase assay

2 µg of Atg2-18 were incubated with 1 µg of Atg1 variants (wild type, kinase-deficient (D211A), or kinase domain only (1-454)) in kinase reaction buffer containing 25 mM MOPS pH 7.5, 1 mM EGTA, 10 mM Na_3_VO_4_, 15 mM MgCl_2_, 280 nM ATP and 1 uCi gamma-^32^P-ATP (SCP-301; Hartmann Analytic) in a total volume of 12 µl. Reactions were incubated for 20 min at 30°C and terminated by addition of 3 µl 6x SDS-PAGE loading buffer, followed by heating at 95°C for 10 min. Samples were resolved by SDS-PAGE. Gels were subsequently dried for 1 h at 80°C, and incorporation of ^32^P was detected using a phosphoimager (GE Healthcare).

### Microscopy-based lipid transfer assay

GUVs were added to a 96-well plate coated with GUV buffer + 5 mg/mL BSA. 200 nM Atg1, 1 mM ATP (where indicated), 2 mM MgCl_2_ and GUV buffer were added. The reaction was started by the addition of 200 nM Atg2-Atg18 or Atg2-Atg18 variants imaged after 5 minutes to allow diffusion and sinking of the GUVs. Images were acquired with spinning disk microscopy, every 30 seconds for 50 minutes at 30°C. Analysis was performed with Fiji.

### Sample preparation for mass spectrometry analysis

Samples for in vitro phosphomapping were prepared as described previously in section “Protein expression and purification”. Atg2-GFP—Atg18 was mixed with Atg1, ATP and MgCl_2_. The control sample did not contain ATP. After incubation the reaction was loaded on a Superose 6 Increase 10/300 column (29091596; Cytiva) and fractions containing Atg2 were collected.

For samples for targeted in vivo phosphomapping yeast cells were grown as explained below (see method section “Nitrogen starvation experiment”). Cells for the “rich” conditions were not in nitrogen starvation medium but harvested directly. For the “starvation” condition cells were harvested after 1 h in starvation medium. Cleared cell lysate was incubated with acetylated GFP-trap beads (Hollenstein *et al*, 2023) for 45 min. Beads were washed and resuspended in 1x PBS.

### Sample processing mass spectrometry analysis

For in vitro phosphomapping samples designated for tryptic digestion, approximately 1.5 µg protein in solution was filled up to 10 µL with 50 mM ammonium bicarbonate (ABC) and then 1 µL of 20 % sodium deoxycholate (SDC) in 50 mM ABC was added. Proteins were reduced by adding 0.44 µL of 250 mM dithiothreitol (DTT) for 30 min at 50°C before adding 0.44 µL of 500 mM iodoacetamide (IAA) and incubating for 30 min at room temperature in the dark. Remaining IAA was quenched with 0.22 µL of 250 mM DTT for 10 min. The solution was diluted to 1% SDC by adding 10 µL 50mM ABC. Proteins were digested with 50 ng trypsin (Trypsin Gold, Promega) at 37°C overnight. The digest was stopped by the addition of 10% trifluoroacetic acid (TFA) to a final concentration of 2%. Then the samples were centrifugated at 18,500 g for 10 min, and the supernatant was transferred to custom-made C18 stagetips for desalting (Rappsilber *et al*, 2007). The eluates were reduced to dryness in a vacuum centrifuge and taken up in 20 µL of 0.1% TFA in 2% acetonitrile (ACN).

For samples designated for dual-protease digestion, approximately 1.8 µg protein in 4 µL solution were mixed with 16 µL of 8 M urea in 50 mM ABC. Proteins were reduced by adding 0.8 µL of 250 mM DTT for 30 min at RT before adding 0.8 µL of 500 mM IAA and incubating for 30 min at room temperature in the dark. Remaining IDAA was quenched with 0.4 µL of 250 mM DTT for 10 min. The solution was diluted to 1 M urea by adding 108 µL of 50 mM ABC. Proteins were digested with 60 ng trypsin (Trypsin Gold, Promega) at 37°C overnight. Thirty % of the digest was kept as “tryptic” digest. The remaining 70% were incubated with 40 ng chymotrypsin (Sequencing Grade, Promega) for 1 h at 25°C (“tryptic/chymotryptic”). The digests were stopped by the addition of 10% TFA to a final concentration of 0.5% and desalted on custom-made C18 stagetips (Rappsilber *et al*, 2007). The eluates were reduced to dryness in a vacuum centrifuge and taken up in 20 µL of 0.1% TFA in 2% ACN.

For targeted in vivo phosphomapping beads were resuspended in 50 µL 1 M urea and 50 mM ammonium bicarbonate. Disulfide bonds were reduced with 2 µL of 250 mM dithiothreitol (DTT) for 30 min at room temperature before adding 2 µL of 500 mM iodoacetamide and incubating for 30 min at room temperature in the dark. Remaining iodoacetamide was quenched with 1 µL of 250 mM DTT for 10min. Proteins were digested with 150 ng LysC (mass spectrometry grade, FUJIFILM Wako chemicals) at 25°C overnight. The supernatant was transferred to a new tube and digested with 150 ng trypsin (Trypsin Gold, Promega) in 1.5µL 50 mM ammonium bicarbonate at 37°C for 5 hours. The digest was stopped by the addition of trifluoroacetic acid (TFA) to a final concentration of 0.5%, and the peptides were desalted using C18 Stagetips (Rappsilber *et al*, 2007).

### Liquid chromatography-mass spectrometry analysis

LC-MS analysis for in vitro phosphomapping was performed on an UltiMate 3000 RSLCnano LC system (Thermo Scientific) coupled to an Orbitrap Exploris 480 mass spectrometer (Thermo Scientific). The system was equipped with a Nanospray Flex ion source (Thermo Scientific), coated emitter tips (PepSep, MSWil), and a Butterfly Portfolio Heater (Phoenix SCT).

Peptides were loaded onto a trap column (PepMap Neo C18 5mm × 300 µm, 5 μm particle size, Thermo Scientific) using 0.1% TFA as mobile phase, and separated on an analytical column (Acclaim PepMap 100 C18 HPLC Column, 50 cm × 75 µm, 2 μm particle size, Thermo Scientific), applying a linear gradient starting with a mobile phase of 98% solvent A (0.1% FA) and 2% solvent B (80% acetonitrile, 0.08% FA), increasing to 35% solvent B over 80 min at a flow rate of 230 nl/min. The analytical column was heated to 30°C.

The mass spectrometer was operated in data-dependent acquisition (DDA) mode, with 2 s MS1 cycle time. Survey scans were acquired from 350-1500 m/z with lock mass enabled, normalized AGC target of 100%, resolution of 60,000. The most intense precursor ions (charge states +2 to +6) were selected for fragmentation using an isolation window of 1.2 m/z. Selected ions were analyzed with a maximum fill time of 100 ms, normalized AGC target of 100%, and resolution of 15,000 after HCD fragmentation with normalized collision energy of 28%. Monoisotopic precursor selection (MIPS) was set to “peptide” mode, the intensity threshold to 1E5, and selected precursors were dynamically excluded for 20 seconds with isotope exclusion enabled.

LC-MS analysis for targeted in vivo phosphomapping was performed on a Vanquish Neo UHPLC system (Thermo Scientific) coupled to an Orbitrap Astral mass spectrometer (Thermo Scientific). The system was equipped with a Nanospray Flex ion source (Thermo Scientific), and a Column Heater (IonOpticks) connected to a Heater Controller (IonOpticks)

Peptides were loaded onto a trap column (PepMap Neo C18 5mm × 300 µm, 5 μm particle size, Thermo Scientific) using 0.1% TFA as mobile phase, and separated on an analytical column (Aurora Ultimate XT C18, 25 cm × 75 µm, 1.7 µm particle size, IonOpticks), applying a linear gradient starting with a mobile phase of 98% solvent A (0.1% FA) and 2% solvent B (80% acetonitrile, 0.08% FA), increasing to 45% solvent B over 30 min at a flow rate of 300 nl/min. The analytical column was heated to 50°C.

The mass spectrometer was operated in data-dependent acquisition (DDA) and Parallel Reaction Monitoring (PRM) mix mode. Survey scans were acquired from 375-1500 m/z, normalized AGC target of 100%, resolution of 120,000. The DDA cycle time is set to 0.3s. The most intense precursor ions (charge states +2 to +6) were selected for fragmentation using an isolation window of 1.6 m/z. Selected ions were analyzed with a maximum fill time of 5 ms, normalized AGC target of 100%, with normalized collision energy of 30%. Monoisotopic precursor selection (MIPS) was set to “peptide” mode, the intensity threshold to 5E3, and selected precursors were dynamically excluded for 15 seconds with isotope exclusion enabled. The unscheduled PRM targets are SIǪDLTNSMLǪFSDPLTTYNR (unmodified, 1222.5917++), <u>S</u>IǪDLTNSMLǪFSDPLTTYNR (single phosphorylated, 1262.5749++), and <u>S</u>IǪDLTN<u>S</u>MLǪFSDPLTTYNR (double phosphorylated, 1302.5581++). Precursor of peptides of interest were isolated in a 0.7 m/z window and fragmented with 27% HCD collision energy. The normalized AGC target to 300%. Maximal injection time for target peptides was set to 50 ms.

### Mass spectrometry data analysis

MS raw data of the in vitro phosphomapping were analyzed with FragPipe (20.0), using MSFragger (3.8) (Kong *et al*, 2017), IonǪuant (1.9.8) (Yu *et al*, 2021), and Philosopher (5.0.0) (Da Veiga Leprevost *et al*, 2020). The default FragPipe workflow for label free quantification (LFǪ-MBR) was used, except “Normalize intensity across runs” was turned off. Cleavage specificity was set to Trypsin/P with two allowed missed cleavages, or to a combination of Trypsin/P and Chymotrypsin (cleavage C-terminal to F/Y/W/L/K/R, except when followed by P) with four allowed missed cleavages. The protein FDR was set to 1%. Carbamidomethyl was used as fixed cysteine modification. Oxidation of methionine, N-terminal protein acetylation, and phosphorylation of serine, threonine, and tyrosine (STY) were specified as variable modifications, with a maximum of 3 variable modifications allowed per peptide. MS2 spectra were searched against the S. cerevisiae 1 protein per gene reference proteome from Uniprot (Proteome ID: UP000002311, release 2022.01), concatenated with a database of 379 common laboratory contaminants (release 2023.01, https://github.com/maxperutzlabs-ms/perutz-ms-contaminants).

Computational analysis was performed using Python and the in-house developed Python library MsReport Python library (version 0.0.16; source code: https://github.com/hollenstein/msreport) (Hollenstein, 2026). LFǪ ion intensities were log2-transformed and normalized to the corresponding log2 LFǪ protein intensity. To analyze phosphorylation at the site level, “site groups” were defined, which represent the unique combinations of co-occurring phosphorylation sites identified on the same peptide sequence. Individual precursor ion intensities were aggregated to their respective site groups using the MaxLFǪ algorithm. Each site group was assigned the maximum isoform localization probability observed among all its constituent ions. For each phosphorylation site group, corresponding “counter-ions” were defined as any non-phosphorylated peptide overlapping with any of the phosphorylated residues of that site group. For each site group, its counter-ions were aggregated using the MaxLFǪ algorithm to generate a corresponding “LFǪ counter intensity”. The Python library XlsxReport (version 0.0.5; source code: https://github.com/hollenstein/xlsxreport) (Hollenstein & Hartl, 2025) was used to create formatted Excel files summarizing the results of the proteomics experiments (Dataset EV1).

PRM measurements from the targeted in vivo phosphomapping were analyzed in Skyline (26.1.1.279, (Pino *et al*, 2020)). After loading the list of the 3 PRM targets, the raw data were imported. Database search in Skyline with MsFragger (Kong *et al*, 2017) was used to build spectra library and match the peptide IDs to the extracted peaks. All peptides and their transitions were validated manually based on retention time, relative ion intensities, and mass accuracy. Three of the most intense non-interfering transitions of the target peptides were selected and their peak areas were exported from Skyline. The summed fragment peak areas were used for peptide quantification.

### AlphaFold3 predictions

Structures of full-length *S. cerevisiae* Atg2 (UniProt P53855, residues 1–1592) were predicted with AlphaFold3 (Bauer *et al*, 2022) on the AlphaFold Server with default settings. Phosphorylated variants were predicted directly by specifying S109 and/or S116 as phosphoserine (CCD code SEP) in the input, i.e., phosphorylation was not introduced after the prediction. For each variant (Atg2^WT^, Atg2^S109ph^, Atg2^S116ph^ and Atg2^S109ph,S116ph^), three predictions with different random seeds were performed, each yielding five ranked models (15 models per variant). The five models (ranks 1–5) of one prediction per variant were used as starting structures of the CG simulations, and residues 1–230 of two of these models as starting structures of the AA simulations.

Per-residue pLDDT values were obtained by averaging the atom pLDDT values of each residue, and the predicted aligned error (PAE) was averaged over all tokens of each residue (phosphoserine is represented by one token per atom). The width of the N-terminal groove was quantified in all 15 models per variant by the distances between Ser40 Oγ and Met117 Cε, Val79 Cγ2 and Ile110 Cγ2, and Leu106 Cδ1 and Ile178 Cγ2 using MDAnalysis (Michaud-Agrawal *et al*, 2011). All AlphaFold3 output files (model coordinates, summary confidences including ranking scores, full data files with pLDDT and PAE, and job requests including seeds) are deposited in the Zenodo repository (DOI: 10.5281/zenodo.23479169).

### All-atom simulations of the N-terminus in solution

Simulations were run using the GROMACS/2022.4 version (Bauer *et al*, 2022). Two starting conformations of the N-terminal region (residues 1–230) were taken from the full-length AlphaFold3 models: a closed conformation (residues 1–230 of the rank-1 Atg2^WT^ model) and an open conformation (residues 1–230 of the rank-2 Atg2^S109ph^ model), in which H1 and H2 are slightly displaced from the groove with respect to the WT model. We chose the rank-2 model due to its lower PAE in the H2 region (Appendix Fig. S2). Each phosphorylation state (Atg2^WT^, Atg2^S109ph^, Atg2^S116ph^ and Atg2^S109ph,S116ph^) was simulated in both conformations. Phosphorylation of S109 and S116 (individually and in combination) was introduced manually using CHARMM-GUI (Jo *et al*, 2008). To avoid artefacts, phosphate groups were modelled in the mono-deprotonated state. All systems were simulated in 150 mM NaCl solution using the charmm36m force field (Huang *et al*, 2017) and TIP3P water, with LINCS to fix bond lengths involving hydrogen atoms. Energy minimization was performed using the steepest descent algorithm for 5,000 steps.

Equilibration was carried out for 12.5 ns with a 1 fs time step, using initial velocities drawn from the Maxwell–Boltzmann distribution for independent replicas. Temperature was maintained at 310 K (37°C) using the Berendsen thermostat with a coupling constant of 1 ps (Berendsen *et al*, 1984). Position restraints of 400 kJ mol⁻¹ nm⁻² were applied to backbone heavy atoms and 40 kJ mol⁻¹ nm⁻² to side chains.

Three independent production simulations of 1 µs were performed per system (24 simulations in total) with a 2 fs time step. Pressure was maintained at 1 bar using the Parrinello–Rahman barostat (Parrinello & Rahman, 1981) applied isotropically with a coupling constant of 5 ps and a compressibility of 4.5 × 10⁻⁵ bar⁻¹. Temperature was controlled using the velocity-rescale thermostat (Bussi *et al*, 2007) at 310 K (37°C) with a coupling constant of 1 ps. Electrostatic interactions were treated using the particle mesh Ewald method (Essmann *et al*, 1995) with a cutoff of 1.2 nm; the same cutoff was applied to van der Waals interactions. A Verlet cutoff scheme was used with a cut-off distance of 1.272 nm for the short-range neighbor list to avoid neighbor list artefacts(Kim *et al*, 2023). For the analysis, all production frames saved every 1 ns were considered.

### Coarse-grained simulations of lipid entry

Simulations were run using the GROMACS/2022.4 version (Bauer *et al*, 2022) with the Martini 3 force field (Souza *et al*, 2021). The five AlphaFold3 models of each variant (Atg2^WT^, Atg2^S109ph^, Atg2^S116ph^ and Atg2^S109ph,S116ph^; full-length Atg2, residues 1–1592) were converted to CG resolution with Martinize2 (Kroon *et al*, 2025) (vermouth 0.9.2), using position restraints in the backbone beads. Secondary structure was assigned with DSSP (Kabsch & Sander, 1983), and an elastic network with a force constant of 700 kJ mol⁻¹ nm⁻² was applied between backbone beads within 0.5–0.9 nm. For each variant, the protein topology was generated from one reference model (rank 1 for Atg2^WT^, Atg2^S109ph,S116ph^ and Atg2^S116ph^, rank 2 for Atg2^S109ph^) and used for all five models of that variant. Elastic-network bonds were removed in flexible regions (residues 1–5, 22–30, 93–102, 127–154, 353–370, 479–489, 614–640, 762–780, 920–1023, 1419–1497, and 1587–1592) and for H1 (residues 6–21) and H2 (residues 104–118), so that the two helices were not restrained to the geometry of the reference model. We used two beads to describe the side chain of phosphoserine: a bead SC1 (type TP1; atoms CB and OG) bonded to the backbone bead (with a bond length 0.287 nm and a force constant 5,000 kJ mol⁻¹ nm⁻²) and a phosphate bead PO4 (type Ǫ5n, charge −1; atoms P, O1P, O2P and O3P) bonded to SC1 (with bong length of 0.400 nm and a force constant of 5,000 kJ mol⁻¹ nm⁻²). This mapping strategy has been previously used by Mehranfar et al. (Mehranfar *et al*, 2026). Phosphoserine was treated as a standard amino acid for secondary-structure assignment.

For each model, ten independent starting configurations were generated by randomly inserting 2,500 POPC lipids around the protein in a 25 × 25 × 25 nm³ simulation box. POPC parameters from the initial Martini 3 release were used (Souza *et al*, 2021). Martini 3 water was added using a van der Waals radius of 0.21 nm, and Na⁺ and Cl⁻ ions were included to neutralize the system and achieve a final concentration of 150 mM. Initial velocities were assigned from the Maxwell–Boltzmann distribution at 310 K (37°C).

Following energy minimization (1,000 steps, steepest descent), ten 1 μs simulations per starting configuration (200 simulations in total) were performed using a 20 fs time step. Temperature was maintained at 310 K (37°C) using the Berendsen thermostat with a coupling constant of 2 ps, and pressure was maintained at 1 bar using the Berendsen barostat applied isotropically with a coupling constant of 6 ps (Berendsen *et al*, 1984). Compressibility was set to 3 × 10⁻⁴ bar⁻¹. Electrostatic interactions and van-der-Waals interactions were cut-off at 1.1 nm. An increased Verlet cutoff scheme was used with a cut-off distance of 1.418 nm for the short-range neighbor list. Every 1 ns of the production run was used for the analysis.

For the analysis, all trajectories were superposed on the backbone beads of a common reference structure, so that the hydrophobic groove is aligned with the z axis. The density of lipid head groups along the groove was calculated from the PO4 beads of POPC within a 30 Å × 30 Å column centred on the protein’s centre of mass (220 Å along z, 1 Å bins), averaged over time for each simulation and then over simulations (mean ± SEM). The displacement of H2 was calculated per frame as the mean distance between the backbone beads of H2 and the Cα atoms of H2 in the respective starting AlphaFold3 model, after superposition of the backbone of the N-terminal core (residues 35–100 and 176–229). For visualization, CG protein structures were backmapped to atomistic resolution with CG2AT (Vickery & Stansfeld, 2021).

### All-atom simulations of membrane insertion

Simulations were run using the GROMACS/2022.4 version (Bauer *et al*, 2022) using the charmm36m force field (Huang *et al*, 2017) TIP3P water, with LINCS to fix bond lengths involving hydrogen atoms. The H1 helix and the N-terminal segment (residues 1–20, including a C-terminal methyl capping group) were embedded using CHARMM-GUI (Jo *et al*, 2008) in a membrane composed of 50% POPC, 20% POPE, 20% POPI, and 10% ergosterol, with 150 mM NaCl to ensure overall charge neutrality. The system was energy-minimized using 5,000 steps of steepest descent.

Equilibration was performed in a stepwise manner, consisting of 2.5 ns in the NVT ensemble followed by 16.25 ns in the NPT ensemble, with gradually decreasing position restraints applied to backbone, side chain and lipid phosphate beads (initial force constants set to 4000, 2000 and 1000 kJ mol⁻¹ nm⁻², respectively) alongside progressively reduced lipid dihedral restraints (initially 1000 kJ mol⁻¹ nm⁻²). After the first 1.25 ns of the NPT phase, the time step was increased from 1 to 2 fs. Temperature was maintained at 310 K (37°C) using the Berendsen thermostat with a coupling constant of 1 ps (Berendsen *et al*, 1984). During NPT equilibration, pressure was maintained at 1 bar using the Berendsen barostat applied semi-isotropically with a coupling constant of 5 ps (Berendsen *et al*, 1984). The compressibility was set to 4.5 × 10⁻⁵ bar⁻¹.

For each of three independent replicas, initial velocities were assigned from the Maxwell–Boltzmann distribution at 310 K (37°C). Production simulations of 1 μs were subsequently performed. Electrostatic interactions were treated using the particle mesh Ewald method (Essmann *et al*, 1995), with a cutoff of 1.2 nm for all real-space interactions. The thermostat and barostat were switched to the velocity-rescale thermostat and Parrinello–Rahman barostat, respectively (Parrinello & Rahman, 1981; Bussi *et al*, 2007). A Verlet cutoff scheme was used with a cut-off distance of 1.373 nm for the short-range neighbor list. For the analysis, all production frames saved every 1 ns were considered. A summary of all simulations performed in this study is provided in Table EV3.

### AlphaFold2 Multimer screening

Pairwise protein-protein interactions were screened using the HT-ColabFold pipeline (Hohmann *et al*, 2026) (https://gitlab.com/BrenneckeLab/ht-colabfold/-/tree/2afda28cf8e1b81228370bcbe75db18bc9253daf/). HT-ColabFold is based on ColabFold (Mirdita *et al*, 2022), which runs AlphaFold2-Multimer (Jumper *et al*, 2021; preprint: Evans *et al*, 2021). Scs2 was screened against 168 *S. cerevisiae* proteins annotated with the UniProt keyword “Autophagy”. Each prediction generates five models, relaxation was omitted. Predictions were ranked both by the maximum interface predicted TM-score (iPTM) and by the maximum PEAKscore across the five models. PEAKscore is a metric provided by the HT-ColabFold pipeline that reflects the minimum inter-chain predicted aligned error (PAE), scaled to a 0-1 range where higher values indicate more confident interfaces. For each hit, predicted structures and associated PAE and pLDDT plots were inspected manually. The pipeline was run on the Life Science Compute Cluster (LISC, University of Vienna).

### Atg9-GFP-vesicle isolation

Atg9-GFP-vesicles were isolated as described by (Sawa-Makarska *et al*, 2020). Yeast cells expressing Atg9-GFP-TEV-TAP (SMY276) were grown using a fermenter (New Brunswick 13 l vessel) and harvested by centrifugation at 2500 x g for 20 min at 4°C. The cell pellet was washed with PBS buffer pH 6.8 containing 2% (w/v) glucose and resuspended in lysis buffer (50 mM NaCl, 25 mM Hepes pH 7.5, 750 mM sorbitol, 5 mM EDTA, cOmplete protease inhibitors (5056489001; Roche), flash frozen as droplets in liquid nitrogen and opened mechanically using a freezer mill. 24 g of freezer milled powder were thawn in HSE buffer containing protease inhibitors (135 mM NaCl, 15 mM Hepes pH7.5, 750 mM sorbitol, cOmplete protease inhibitors (5056489001; Roche), an FY-inhibitor mix (39104; Serva)) and cleared by centrifugation at 146900 x g at 4°C for 25 min (Beckman Coulter XPN-90, rotor Ti45). The supernatant was incubated with 200 µl IgG-coupled (I5006, Sigma-Aldrich) M-270 epoxy dynabeads (14302D; Invitrogen) overnight at 4°C. The beads were washed once each with HSE buffer containing protease inhibitors, HSE high salt buffer (250 mM NaCl, 15 mM Hepes pH 7.5, 750 mM sorbitol), HSE buffer (135 mM NaCl, 15 mM Hepes pH 7.5, 750 mM sorbitol), and HSE buffer without sorbitol (135 mM NaCl, 15 mM Hepes pH 7.5). The beads were resuspended in 400 µL buffer (150 mM NaCl, 30 mM Hepes pH 7.5) and final Atg9-GFP-vesicles were cleaved with TEV protease for 3 h at 4°C.

### Microscopy-based pull-down

Scs2^MSP^-Halo-GST was bound to GST-Trap agarose beads (17075601; Cytiva) or Halo-Trap agarose beads (ota-20; Chromotek) for 1 h at 4°C. Beads were washed in reaction buffer (150 mM NaCl, 25 mM Hepes pH 7.5, 1mM DTT; for Fig. 6D the buffer contained 150 mM NaCl, 30 mM Hepes pH7.5) and prey proteins were added at a final concentration of 0.5 µM. 0.5 mM MgCl_2_ and, where indicated, 0.1 mM ATP were added. Prey proteins were incubated at room temperature away from light for 15 min prior imaging. For Fig. 6D the beads were incubated with the prey proteins for 25 min at room temperature and subsequently, 65 µL of isolated Atg9-vesicles were added and incubated at 4°C overnight. Beads were imaged at room temperature with a Plan-Apochromat 20×/0.8 WD 0.55 mm objective on a Zeiss LSM700 confocal microscope and the Zeiss ZEN 2012 SP5 imaging software running on Windows 10 (64-bit). Signal intensities were quantified with the aid of a machine learning tool as previously reported (Ferrari *et al*, 2024).

### Yeast strains and media

Strains for overexpression and subsequent purification of Atg2-18 variants derive from genetically modified *S. cerevisiae* BY4741. Strains for *in vivo* analyses derive from genetically modified *S. cerevisiae* w303. Generated strains are listed in Table EV1. Knockouts, tags and other modifications were obtained via homologous recombination of PCR products of interest. Briefly, log-phase cells were washed with lithium acetate (100 mM) and incubated in a mix containing lithium acetate (L6883; Sigma-Aldrich), PEG 2000 (202509; Sigma-Aldrich), salmon sperm DNA (15632011; Invitrogen), and the PCR product of interest at 30°C for 30 min and then shifted to 42°C for 20 min.

### Nitrogen starvation experiments

Cells were grown in synthetic medium (0.17% (wt/vol) yeast nitrogen base without amino acids and ammonium sulfate (233520; BD), 0.5% (wt/vol) ammonium sulfate (31119-M; Sigma-Aldrich), 2% (wt/vol) a-D-glucose monohydrate (16301; Sigma-Aldrich), supplemented with complete or selective amino acid mix for plasmid selection. Log-phase cells were harvested, washed three times, and resuspended in SD-N medium (0.17% (wt/vol) yeast nitrogen base without amino acids and ammonium sulfate (233520; BD) and 2% (wt/vol) a-D-glucose monohydrate (16301; Sigma-Aldrich)). Cells were then incubated at 30°C shaking and imaged at defined time points.

### Giant ApeI assay

Cells expressing pRS425-prCUP1-APE1-prAPE1-tagBFP-APE1 plasmid were grown to logarithmic phase and incubated for 3h in selective medium supplemented with 250μM of CuSO4 (UN3288; VWR) Cells were subsequently washed three times and shifted to nitrogen starvation medium. Imaging was performed every minute, 46 times, starting after 45 min of starvation at 30°C. Images were acquired using spinning disk microscopy. Analysis was performed as previously reported (Schütter *et al*, 2020).

### Autophagosome biogenesis duration

Cells were starved for nitrogen (see above). Imaging was performed every minute, 46 times, starting after 45 min of starvation at 30°C. Images were acquired using spinning disk microscopy. Analysis was performed as previously reported (Dabrowski *et al*, 2023).

### Fluorescence microscopy

Cells were imaged at room temperature in a 96-well plate with a glass bottom (655892; Greiner Bio-One) using an inverted microscope Zeiss Axio Observer Z1 equipped with, an EC Plan-Neofluar 100×/1.45 Oil, an Orca Flash 4.0 LT+ camera, and a ZEN blue 3.3 pro software on Windows 11 pro 64-bit. Alternatively, experiments were performed with an inverted Nikon Ti2-E microscope (RRID:SCR_021068) equipped with a Yokogawa CSU-rX1-A1 Nipkow spinning disk, a CFI Plan Apo λ 100×/1.45 Oil objective, a EM-CCD back-rilluminated Andor iXon Life 888, and a VisiView 7.0 (Visitron Systems) software running on Windows 10 (64 bit). Images were acquired at indicated time points and deconvolved with Huygens Professional 24.10 (Scientific Volume Imaging) and analyzed using Fiji Version 2.16.0/1.54p (RRID:SCR_002285).

### Western blot and autophagic flux analysis

0.5 OD600 yeast cells were collected and lysed with 0.255 M NaOH (S492060; Thermo Fisher Scientific). Proteins were precipitated with 100% trichloroacetic acid (T0699; Sigma-Aldrich) and washed once with cold acetone (CP40.3; Roth). Protein pellets were resuspended in protein loading buffer and analyzed by SDS-PAGE using mouse anti-GFP antibody (Max Perutz Labs, Monoclonal antibody facility) in 3% (wt/vol) milk powder (T145; Roth) TBST-T (P7949; Sigma-Aldrich). Secondary antibody incubation was performed using Dylight 800 α-mouse (610145002; Rockland Immunochemicals) or Dylight 680 α-rabbit (5366P; Cell Signaling). Detection and analysis were performed using the Li-COR Odyssey CLx fluorescence imager (RRID:SCR_014579; LI-COR Biosciences) and the Image Studio (RRID:SCR_015795, Version 2.1) software. Autophagic flux was assessed as the ratio between the free GFP and the total GFP signal (free GFP and GFP-rAtg8 combined).

### Statistics and Reproducibility

Statistical analyses were performed using GraphPad Prism software 11.

The number of independent experiments is stated in the figure legends. When pair comparison was assessed, two-tailed, unpaired *t* test was performed to assess statistical significance. One-way ANOVA with multiple comparison was used when more than two sets of data were compared. When technical as well as biological replicates were generated, nested statistical analysis was carried out. Error bars indicate either SD or SEM according to the experiment shown and are indicated in the figure legends. Only statistically significant comparisons are indicated in graphs.

## Data and material availability

The mass spectrometry proteomics data have been deposited to the ProteomeXchange Consortium via the PRIDE (Perez-Riverol *et al*, 2022) partner repository with the data set identifier PXD078404 (https://www.ebi.ac.uk/pride/archive/projects/PXD078404) and PXD086119 (https://www.ebi.ac.uk/pride/archive/projects/PXD086119). Raw trajectory data from molecular dynamics simulations, parameter files, and starting structures, are deposited in a Zenodo repository (https://doi.org/10.5281/zenodo.23479169). The source data are collected in a Zenodo repository (https://doi.org/10.5281/zenodo.23723628, https://doi.org/10.5281/zenodo.23982019, https://doi.org/10.5281/zenodo.23982854).

## Acknowledgements

We thank current and former members of the Martens lab and especially Martin Spitzer and Nicolas Coudevylle for help and advice. We also would like to thank Martin Greaf for providing the pRS425-*prCUP1-APE1-prAPE1-tagBFP-APE1* plasmid. We thank the Max Perutz Labs BioOptics. We thank the Max Perutz Labs Mass Spectrometry facility for technical support. Proteomics analyses were performed using the VBCF instrument pool. The AlphaFold screen has been performed using the Life Science Compute Cluster (LiSC) of the University of Vienna. Molecular graphics and analyses in Fig. 2A, Fig. EV2A, Fig. 5D and Fig. EV4E were performed with UCSF ChimeraX., developed at the University of California. A.C.C. and G.H. thank the Max Planck Computing and Data Facility (MPCDF) for computational support. This work was supported by the Austrian Science Fund (FWF) (grant number: 10.55776/P35061) and the Max Planck Society.

## Expanded View Figure Legends

**Figure EV1.**
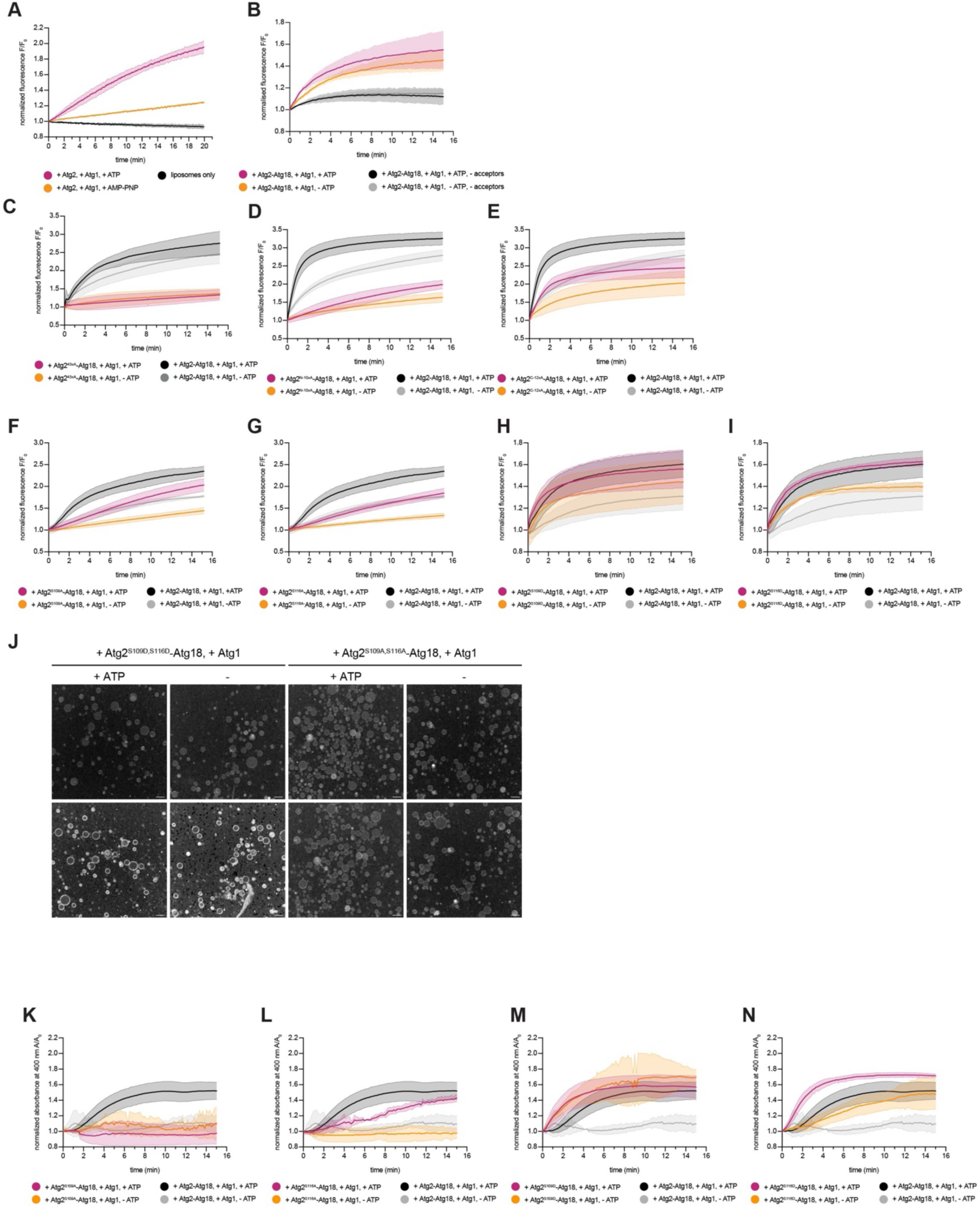
Lipid transfer of Atg2 phospho-mutants is impaired. (**A**) FRET-based lipid transfer assay showing that transfer of Atg2 in the absence of Atg18 is boosted in the presence of active Atg1. (**B**) A slight increase in dequenching by Atg2-Atg18 in the absence of uncolored acceptor liposomes could be observed in the FRET-based lipid transfer assay. (**C**) Transfer by Atg2^43xA^-Atg18 is decreased as measured with a FRET-based lipid transfer assay. (**D-I**) FRET-based lipid transfer assay of Atg2^N-10xA^-Atg18 (**D**), Atg2^C-12xA^-Atg18 (**E**), Atg2^S109A^-Atg18 (**F**), Atg2^S116A^-Atg18 (**G**), Atg2^S109D^-Atg18 (**H**), Atg2^S116D^-Atg18 (**I**). (**J**) Representative images of microscopy-based lipid transfer assay with GUVs and liposomes of indicated Atg2 mutants. Pictures show ATTO 425/ATTO 540Ǫ ratio signal after 0 and 50 minutes of acquisition. Scale bars, 100 µm, quantifications in Fig. 2E,F. (**K-N**) Absorbance-based turbidity assay to test membrane tethering activity of Atg2^S109A^-Atg18 (**K**), Atg2^S116A^-Atg18 (**L**), Atg2^S109D^-Atg18 (**M**), Atg2^S116D^-Atg18 (**N**). All data are means ± SD (n = 3).

**Figure EV2.**
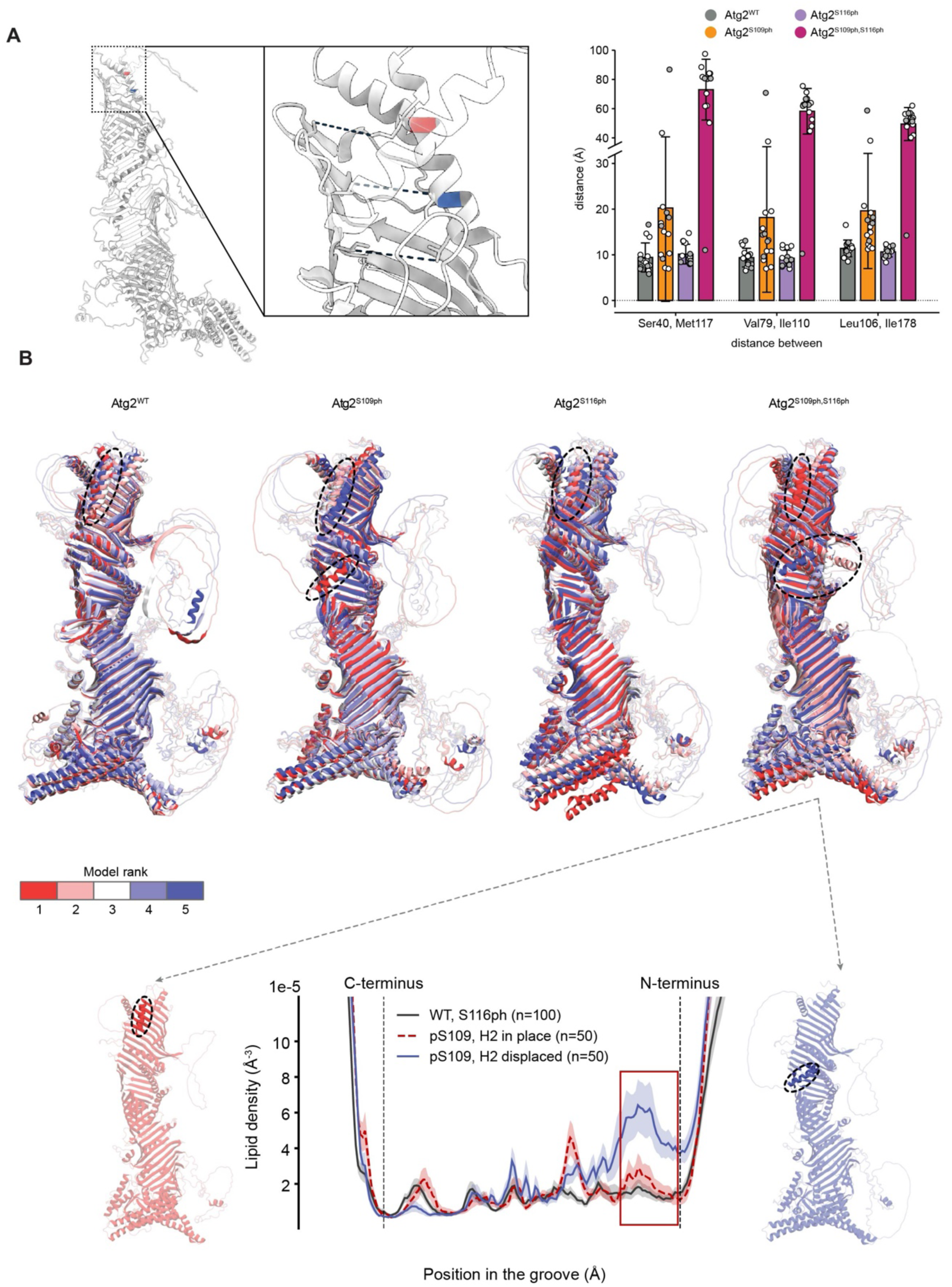
Lipid access in CG simulations depends on the position of H2 in the AlphaFold3 starting models. **(A)** Left: AlphaFold3-predicted structure of Atg2 with S109 (blue) and S116 (red) highlighted (left). Right: Width of the N-terminal groove in the AlphaFold3 models of Atg2^WT^, Atg2^S109ph^, Atg2^S116ph^ and Atg2^S109ph,S116ph^. Distances between Ser40 Oγ and Met117 Cε, Val79 Cγ2 and Ile110 Cγ2, and Leu106 Cδ1 and Ile178 Cγ2, were calculated from the atomic coordinates of the AlphaFold3 model files using MDAnalysis (Michaud-Agrawal et al, 2011). Bars show means ± SD of 15 models per variant (three predictions with different seeds, five ranked models each). Circles show individual models, with the five models used as starting structures for the simulations filled in gray. Note the break in the y-axis. (**B**) AlphaFold3 models of Atg2^WT^, Atg2^S109ph^, Atg2^S116ph^, and Atg2^S109ph,S116ph^. Per variant, five models (ranks 1–5) were used as starting points for CG simulations with dispersed POPC lipids in solution. The degree of displacement of H2 (residues 104–118, indicated with dashed ellipses) varies between the models, as shown by superimposition (top). One model of Atg2^S109ph^ (rank 1) and four models of Atg2^S109ph,S116ph^ (ranks 2–5) show a strong H2 displacement. Bottom: density of lipid head groups along the hydrophobic groove, with the simulations grouped by starting model: Atg2^WT^ and Atg2^S116ph^ (all models, n = 100), S109-phosphorylated variants started from models with H2 in place (Atg2^S109ph^ ranks 2–5 and Atg2^S109ph,S116ph^ rank 1, n = 50), and S109-phosphorylated variants started from models with H2 already displaced in the prediction (Atg2^S109ph^ rank 1 and Atg2^S109ph,S116ph^ ranks 2–5, n = 50). Curves are means of n simulations (1 µs each), with shaded regions indicating SEM. Density calculation and red box as in Fig. 3A.

**Figure EV3.**
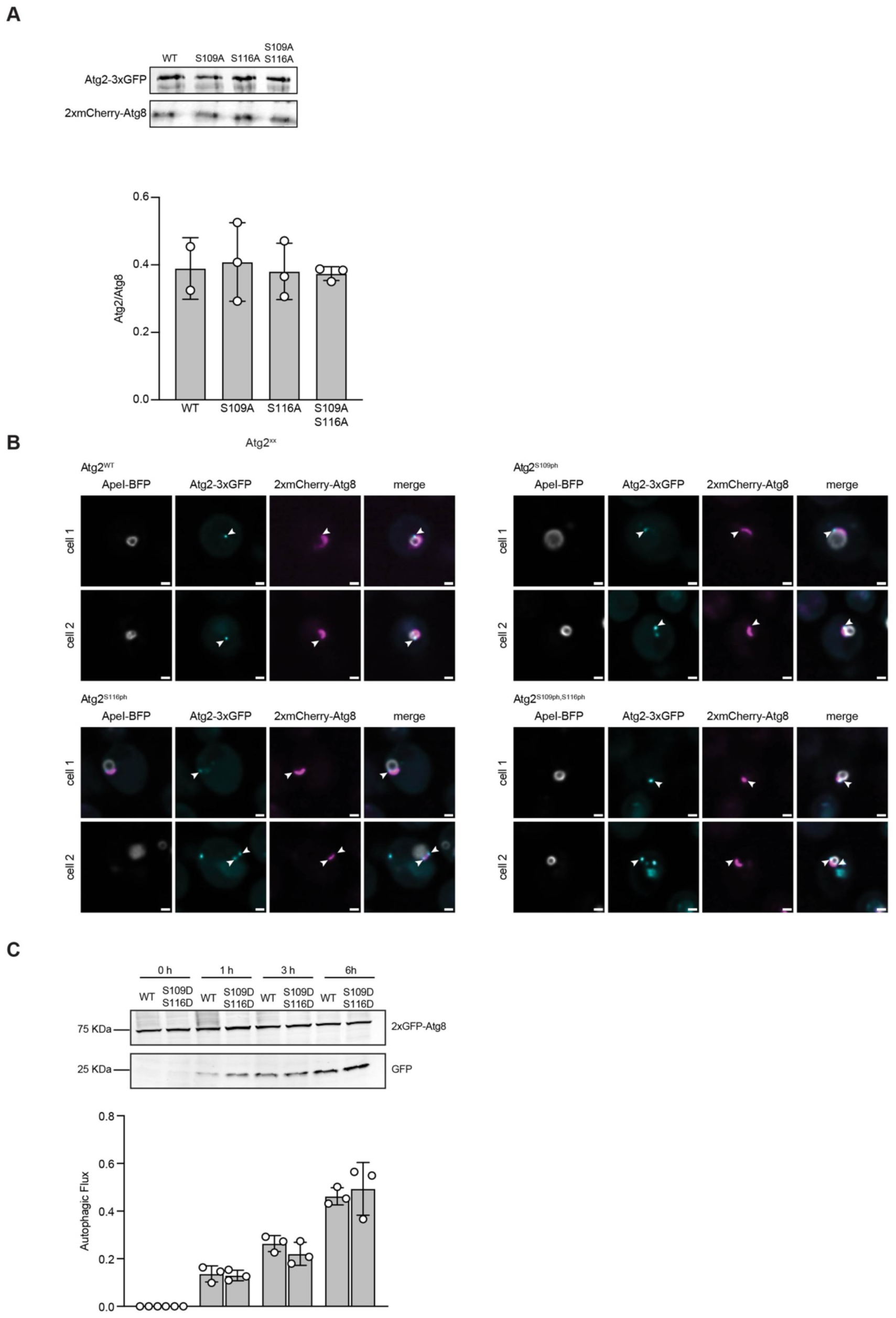
Protein expression levels and localization of the Atg2-variants are unchanged. (**A**) Western blot showing the Atg2-3xGFP expression levels for the indicated variants and its quantification of respective protein levels. Atg2^xx^-3xGFP levels were normalized over 2xmCherry-Atg8 levels. Samples were collected from non-starved cells. Data are means ± SD (n = 3) (**B**) Representative pictures of live spinning disk microscopy of cells expressing Atg2^xx^-3xGFP, 2xmCherry-Atg8 and overexpressing the copper-inducible BFP-preApeI for 3 h. Cells were then shifted to starvation to induce autophagy and imaged after 1 h. Atg2^WT^, Atg2^S109A^, Atg2^S116A^, and Atg2^S109A,S116A^ equally localize to the tip of the phagophore as indicated with white arrowheads. Scale bars: 1 µm. (**C**) Autophagic flux of indicated strains expressing 2xGFP-Atg8 after 0, 1, 3, and 6 h of starvation. Data are means ± SD (n = 3).

**Figure EV4.**
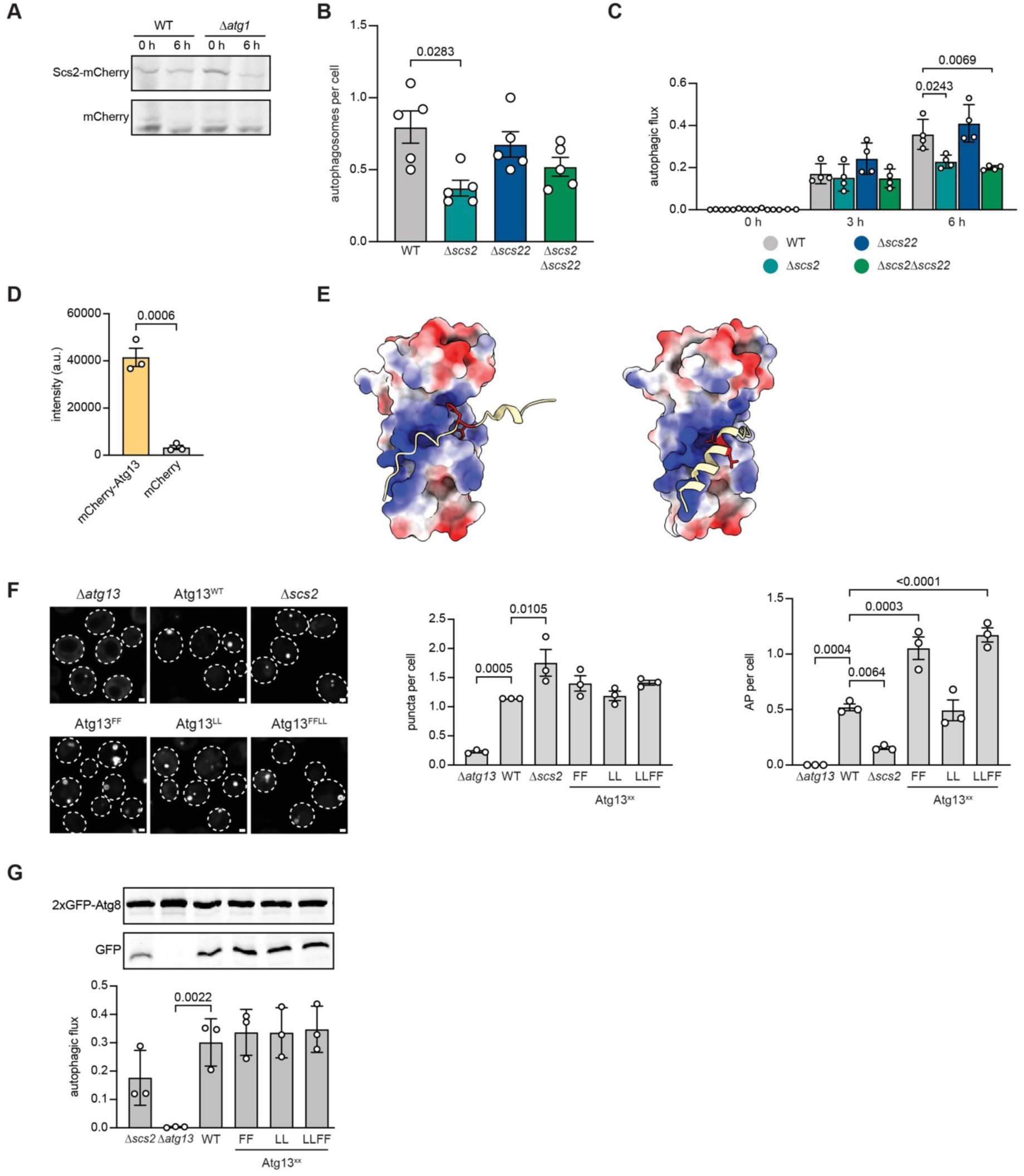
Interaction between the VAP protein Scs2 and Atg13 is important for autophagy. (**A**) Representative western blot analysis of Scs2 levels in growing and starving conditions. No free mCherry signal was detectable independently of treatment or autophagy blockage (n = 3). (**B**) Autophagosome quantification of indicated strains. Live widefield microscopy of cells expressing 2xGFP-Atg8 during nitrogen starvation after 55 min. Data are means ± SEM (n = 5, 50 cells each). Nested One-way ANOVA. (**C**) Autophagic flux of indicated strains expressing 2xGFP-Atg8 after 0, 3, and 6 h of starvation. Data are means ± SD (n = 4), nested One-way ANOVA vs WT of each time point. (**D**) Ǫuantification of Atg13 recruitment to Scs2^MSP^-Halo-GST immobilized on beads (Fig. 5E). Data are means ± SEM (n = 3), nested One-way ANOVA. (**E**) AlphaFold predictions of Scs2 MSP domain with Atg13 FFAT motives containing peptides (FF: 717-738 (left), LL: 620-638 (right)). The electrostatic surface of Scs2 MSP domain is displayed with positively charged residues in blue and negatively charged residues in red. FF and LL residues are marked in red. (**F**) Live widefield microscopy of indicated cells expressing 2xGFP-Atg8 during nitrogen starvation (55 min) of Atg8 positive puncta and autophagosomes. Data are means ± SEM (n = 3, 50 cells each), nested One-way ANOVA vs Atg13^WT^. (**G**) Western blot analysis of autophagic flux of indicated strains expressing 2xGFP-Atg8 after 6 h of starvation. Data are means ± SD (n = 3), nested One-way ANOVA vs Atg13^WT^.

**Figure EV5.**
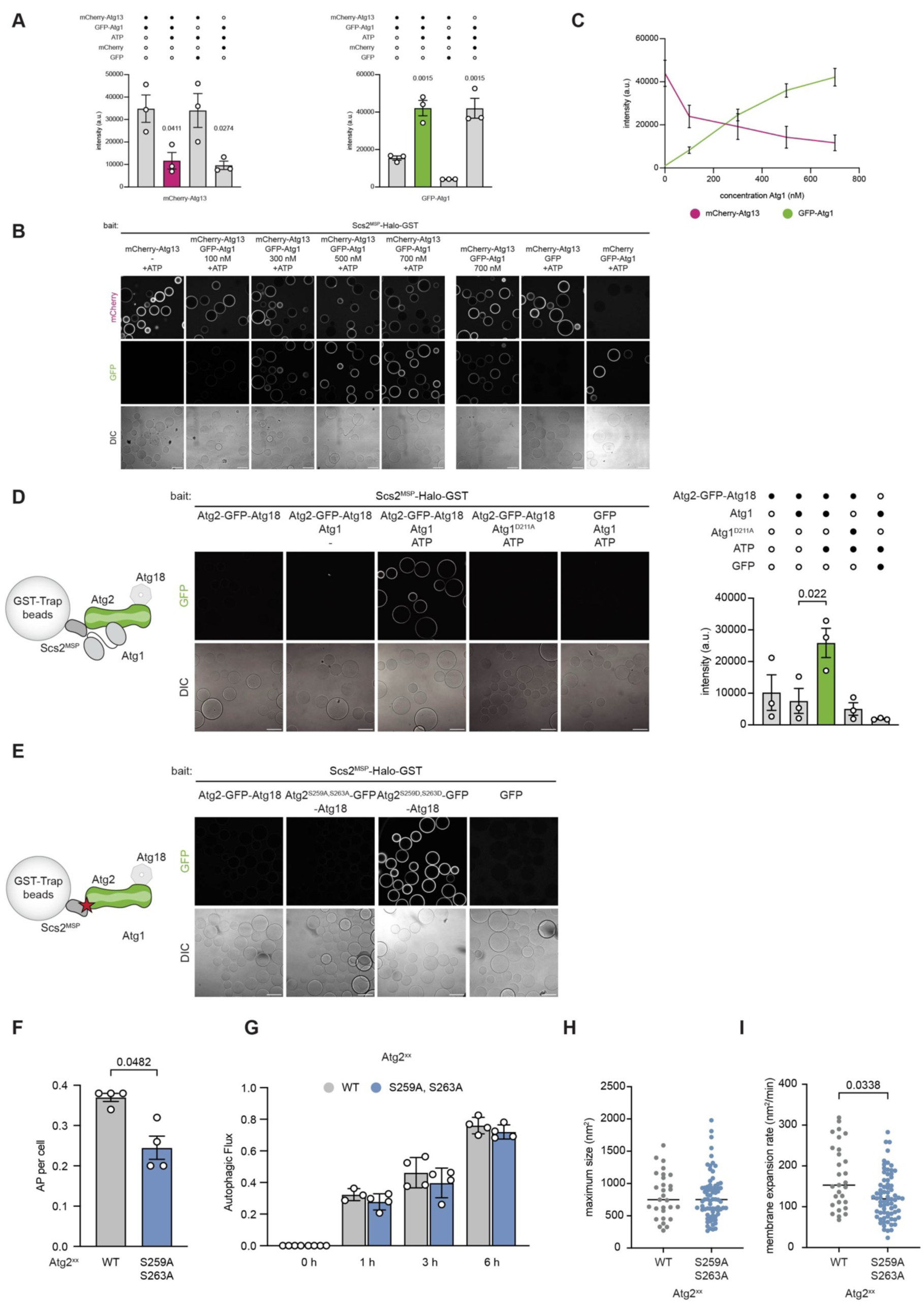
Atg1 competes with Atg13 for binding to Scs2. (**A**) Ǫuantification including negative controls of microscopy-based pull-down assay shown in Fig. 6A. Data are means ± SEM (n = 3), nested One-way ANOVA vs “- ATP” condition. (**B**) Microscopy-based pull-down assay where purified Scs2^MSP^-Halo-GST was immobilized on beads as bait and incubated with mCherry-Atg13 and increasing concentration of GFP-Atg1 in presence or absence of ATP. (**C**) Ǫuantification of mCherry-Atg13 and GFP-Atg1 signal from (**B**) at different GFP-Atg1 concentrations. Data are means ± SEM (n = 3). Fig. 6A and Fig. EV5B were generated analyzing the same dataset and representative pictures are partially repeated. (**D**) Microscopy-based pull-down assay where purified Scs2^MSP^-Halo-GST was immobilized on beads as bait and incubated with Atg2-GFP in presence or absence of Atg1, Atg1^D211A^ and ATP. Data are means ± SEM (n = 3), nested One-way ANOVA vs Atg2-GFP—Atg18 Atg1. (**E**) Microscopy-based pull-down assay where purified Scs2^MSP^-Halo-GST was immobilized on beads as bait and incubated with wild type Atg2-GFP, Atg2^S259A,S263A^-GFP or Atg2^S259D,S263D^-GFP or GFP. Scale bars: 100 µm. (**F**) Ǫuantification of Atg8 positive autophagosome of cells expressing Atg2^WT^ or Atg2^S259A,S263A^ and 2xGFP-Atg8 during nitrogen starvation (1h) Data are means ± SEM (n = 4, 50 cells each), nested t-test analysis. (**G**) Autophagic flux of indicated strains expressing 2xGFP-Atg8 after 0, 1, 3, and 6 h of starvation. Data are means ± SD (n = 4). (**H,I**) Ǫuantification of maximal phagophore expansion (**H**) and membrane expansion rate (**I**) of indicated strains expressing 2xGFP-Atg8 and overexpressing the copper-inducible BFP-preApeI. Nested t-test, n = 4 (median shown).

